# Contractile Centrin Filaments Mediate Mechanical Coupling of Cilia at the Basal Body

**DOI:** 10.64898/2026.11.20.768115

**Authors:** Aparna Sudhakar, Aikaterini M. Kourkoulou, Daphne M. Laan, Yangguang Tian, Gizem Demirkiran, Guillermina R. Ramirez-San Juan

## Abstract

Centrin is a conserved protein that localizes to centrioles and basal bodies across eukaryotes, where it performs scaffolding and structural functions. In ciliates, centrin additionally assembles into patterned, cellular-scale networks required for ultrafast changes in cell shape. Across distantly related algae, centrin forms fibrillar structures that connect the basal bodies to one another and to the nucleus. These features suggest that centrin has cytoskeletal functions, generating and resisting mechanical forces, yet its functions in this context remain poorly understood. Here, we use expansion microscopy and centrin mutants with different network morphologies to study the function of centrin filaments in *Chlamydomonas reinhardtii*. Consistent with previous studies, centrin localizes to the basal bodies and transition zones, to the distal striated fiber, which connects the distal ends of the two basal bodies, and to filaments that extend from the basal bodies toward the nucleus, where they branch into a network that surrounds it. Using mutants of basal body proteins, we show that these centrin fibers assemble independently of basal bodies, which anchor and orient them. Centrin mutants have shorter cilia that regenerate more slowly, consistent with centrin’s structural roles at the basal bodies and transition zones. Beyond these structural roles, centrin serves a mechanical function at the distal striated fiber. Mutants with structural defects in this centrin network show altered ciliary beat frequency and lack synchrony between the two cilia, resulting in impaired swimming. Moreover, the transition zone, the distal striated fiber, and the nucleus–basal body connectors each contract in response to calcium. Together, these results show that centrin assembles into basal body-associated networks that combine structural scaffolding with calcium-responsive contractility and force transmission. Our findings establish centrin as a *bona fide* cytoskeletal protein and provide insight into how a single protein can diversify to perform distinct structural and mechanical functions within the same cell.

---

Centrins are widely conserved proteins, members of the EF-hand family of Ca^2+^-binding proteins [1, 2]. Across eukaryotes they are found associated with centrosomes and basal bodies [3–6]. The function of centrin has been studied mainly in this context, where it contributes to the assembly, maturation, separation and stability of these structures through largely unknown mechanisms [7–10]. At centrioles, centrin is required for structural integrity through its association with Sfi1 [10, 11]. In ciliated cells, centrin is required for ciliogenesis and for the correct duplication and positioning of basal bodies [7, 8, 12, 13]. In yeast, centrin plays an analogous role, as it is essential for duplication of the spindle pole body, the functional equivalent of the centrosome [14].

In protists, in addition to its basal body localization, centrin assembles into higher order filamentous structures and networks with stereotypical geometries. In ciliates, it forms cellular-scale meshes required for the ultrafast shape changes these cells undergo in response to mechanical or chemical stimuli [15–17]. In algae, centrin localizes to fibrillar structures that connect the basal bodies to each other and to the nucleus. Such filamentous networks have been described in both chlorophyte green algae and chrysomonads, suggesting that these centrin-containing structures are conserved across distantly related lineages, however their function remains unknown [18–21].

In *Chlamydomonas*, centrin has been shown to localize to multiple cilia-associated structures, including the ciliary axoneme, transition zone, basal body lumen, the distal striated fiber (DSF) and the nucleus-basal body connectors (NBBCs) [19, 22–26]. Previous studies on centrin mutants and in centrin-depleted cells suggested that this protein is essential for basal body duplication and segregation during cell division [7, 27]. However, the functions of the different fibrillar centrin structures associated with basal bodies remain unclear. Contemporary tools such as expansion microscopy (ExM) and quantitative image analysis now enable the study of the non-canonical functions of centrin in *Chlamydomonas*.

Here, we use these approaches to investigate the cytoskeletal functions of centrin in *Chlamydomonas reinhardtii* and show that centrin performs both structural and mechanical functions at the ciliary base. Consistent with previous studies, we find that centrin localizes to the ciliary transition zone, the basal bodies, the DSF that connects them, and the NBBCs, which extend toward the nucleus and branch into a network of filaments that envelops it. Beyond localization, we examined how these structures are organized in cells with mutations in basal body-associated proteins and in centrin itself. In mutants that lack basal bodies (*bld10*), fibers reminiscent of the NBBCs and the branched network around the nucleus are able to assemble, whereas in mutants with mispositioned basal bodies (*vfl3* and *asq2*), centrin still localizes to the basal bodies, the transition zones, and the

NBBCs, but is absent from the DSF or reduced to a stub. To test how centrin itself contributes to these structures, we took advantage of the previously isolated centrin mutant *vfl2*, in which a single point mutation (E102K) restricts centrin localization to the basal bodies, and of two uncharacterized *vfl2*-derived mutants that carry an additional point mutation (R97H or K105Q) and assemble partial centrin networks. These centrin mutants have defects in ciliary number, length, and regeneration, consistent with centrin’s localization at the basal body and transition zone. These mutants also have structurally abnormal distal fibers and swimming defects that stem from aperiodic ciliary beating or impaired coordination between the two cilia. Finally, we find that the transition zone, DSF, and NBBCs each contract in response to calcium, and that this contraction is selectively impaired in the *vfl2*-derived mutants. Together, these results show that centrin assembles into specialized networks associated with basal bodies that integrate calcium-responsive contractility, force transmission, and structural scaffolding at the ciliary base. Our findings establish that centrin performs cytoskeletal functions at the ciliary base and highlight how a single protein can form assemblies with distinct structural and mechanical functions within the same cell.

## CENTRIN ASSOCIATES WITH BASAL BODIES AND FORMS NETWORKS WITH DISTINCT ARCHITECTURES

In chlorophyte algae, centrin is the main structural component of the fibrous network that connects the basal bodies to each other and to the nucleus (Extended Data Fig. S1)[18, 19]. To study the organization of this network in its cellular context, we used ExM to image centrin in two algae, *Tetraselmis striata*, in which centrin was first identified [18] and *Chlamydomonas reinhardtii* (Fig. 1A,B; see Methods A 3). We co-stained for polyglutamylated tubulin (PolyE), which labels the cilia axoneme, basal bodies and rootlet microtubules, and to visualize the total protein content we stained with NHS-ester. Although the geometry of the fiber array differs between the two, in both species, centrin localizes to the basal bodies and forms fibers that descend from the basal bodies toward the nucleus (Fig. 1A,B).

**FIG 1.**
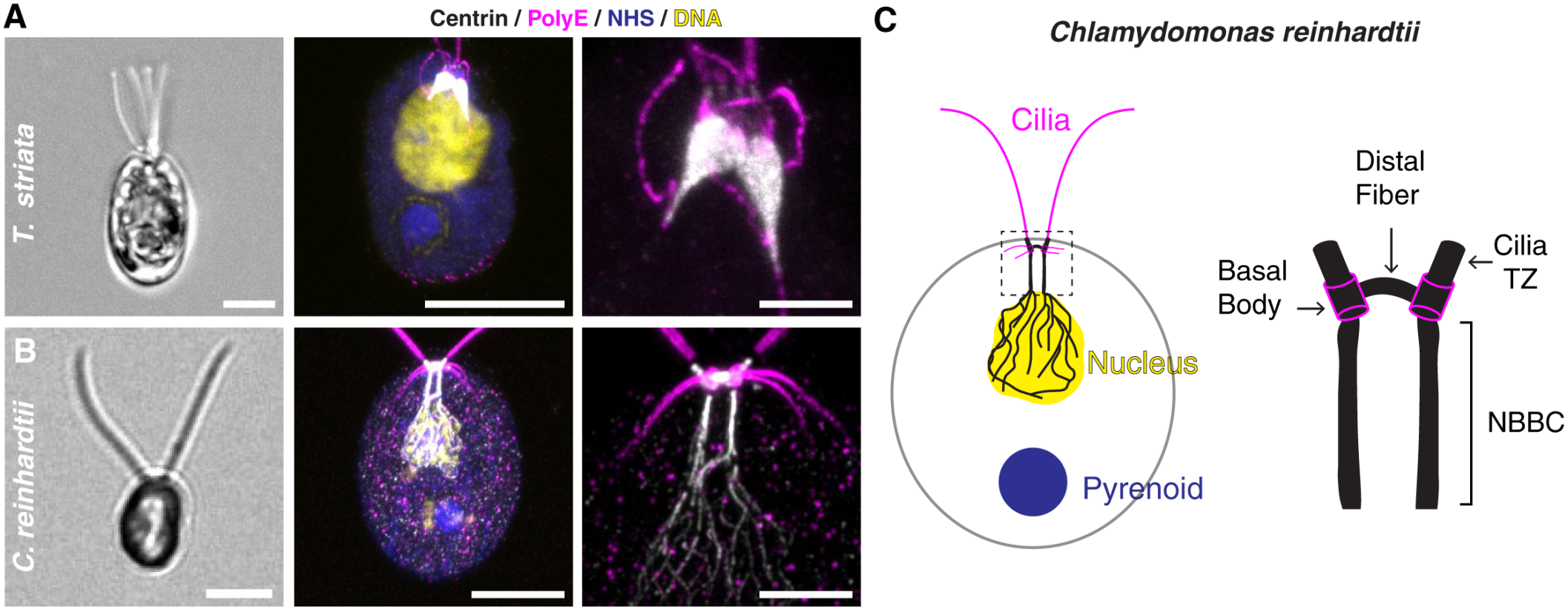
Centrin forms a network that connects distinct basal body-associated structures. (A) Left: brightfield image of *Tetraselmis striata*. Middle: immunofluorescence image stained for centrin (white), PolyE (magenta), NHS-ester (blue) and DNA (yellow), as visualized by expansion microscopy(ExM). Right: inset showing the centrin network. Scale bars: 5 µm (A,B, left and middle), 1 µm (A,B, right). (B) As in (A), for *Chlamydomonas reinhardtii*. (C) Schematic of a *Chlamydomonas* cell showing the substructures to which centrin localizes: the transition zone (TZ), the basal body, the distal fiber connecting the two basal bodies, and the nucleus–basal body connectors (NBBCs), which transition into a branched network around the nucleus.

In *Chlamydomonas*, centrin localizes to distinct sub-structures connected to the basal bodies (Fig. 1B, inset). At the base of each cilium, it is present at the basal bodies and ciliary transition zone. Between the two basal bodies, centrin labels the DSF, which connects them. From the basal bodies, centrin extends toward the nucleus in the form of a fibrillar network that corresponds to the NBBCs. Upon reaching the nucleus, these fibrils transition into branches that envelop the nucleus (Fig. 1C).

These localizations are consistent with previous studies in *Chlamydomonas* [19, 23–26]. These observations show that, within a single cell, centrin adopts diverse architectures ranging from compact accumulations at the basal bodies and transition zone, to fibrils forming the NBBCs, to a branched network around the nucleus. Because all of these structures are organized around the basal bodies, we next asked whether their assembly is directed by the basal bodies.

### BASAL BODIES ORIENT CENTRIN FIBERS BUT ARE DISPENSABLE FOR THEIR ASSEMBLY

Perturbations of centrin impair basal body duplication and segregation [7, 27], but whether the assembly of distinct parts of the centrin network depends on basal bodies remains unclear. To address this, we imaged the centrin networks by ExM in previously characterized *Chlamy-domonas* mutants with defects in basal body structure, positioning, or assembly (Fig. 2A–F).

**FIG 2.**
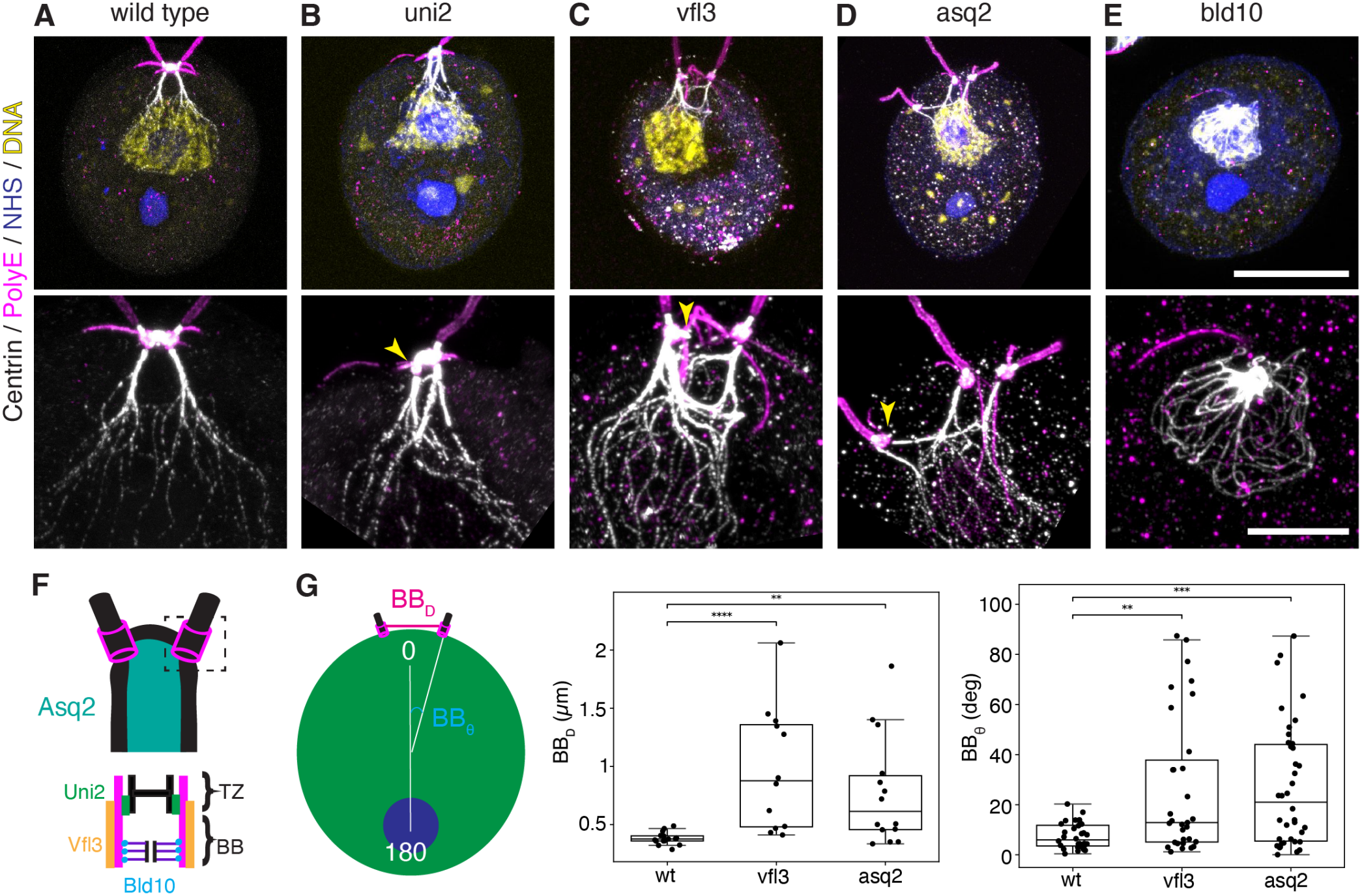
Basal bodies are required for spatial organization of centrin fibers, but not for their assembly. (A) Top: wild-type *Chlamydomonas* cell imaged by ExM and stained for centrin (white), polyglutamylated tubulin (PolyE, magenta), which marks the basal bodies and cilia, total protein (NHS-ester, blue) and DNA (yellow). Bottom: inset showing the centrin network descending from the basal bodies. (B–E) As in (A), for *uni2* (B), *vfl3* (C), *asq2* (D) and *bld10* (E) mutants. Yellow arrowheads indicate the transition zone lacking centrin in *uni2* cells (B) and the DSF stubs in *vfl3* (C) and *asq2* (D) cells. (F) Schematic of the basal body apparatus indicating the localization of Uni2 (green; distal end of the basal body and TZ), Vfl3 (orange; outer wall of the basal body), Bld10 (cyan; cartwheel) and Asq2 (teal; domain between the two NBBCs). Centrin is shown in black and the basal bodies in magenta. (G) Left: schematic defining BB_*D*_, the distance between the two basal bodies, and BB_*θ*_, the angle between the cell axis (defined by the centroids of the cell body and the pyrenoid) and the line connecting the cell-body centroid to the basal body. Middle: BB_*D*_ for wild-type, *vfl3* and *asq2* cells. Right: BB_*θ*_ for the same genotypes. Sample sizes: wild type (CC-124), *n*_cells_ = 16; *vfl3, n*_cells_ = 13; *asq2, n*_cells_ = 15. Boxes denote the 25th and 75th percentiles, central lines the median, and whiskers the most extreme values within 1.5× the interquartile range (IQR). Each mutant was compared with wild type using a two-sided Mann–Whitney *U* test; \*\**p <* 0.01, \*\*\**p <* 0.001, \*\*\*\**p <* 0.0001. Scale bars: 5 µm (A-E, top), 2 µm (A-E, bottom).

We first examined mutants that fail to assemble basal bodies (*bld10* and *bld12*) [28–30]. In these cells, the branched centrin network that surrounds the nucleus remains intact, but the linear fibers that resemble the NBBCs are collapsed or absent (Fig. 2E, Extended Data Fig. S2A). These fibers are extended only in the occasional *bld12* cells with a basal body remnant, in which they run from the nucleus to the remnant (Extended Data Fig. S2A), as previously observed in the mutant *bald-2* [31]. Next we examined how the centrin networks are organized when basal bodies assemble but are abnormal in number and position, as in *vfl3* and *asq2* mutants [32–34]. The basal bodies of these cells appear widely separated (Fig. 2C,D). To quantify this mispositioning, we measured the distance between the two basal bodies and the angle of each basal body relative to the cell-pyrenoid axis (Fig. 2G). Both were significantly larger in the two mutants than in wild-type cells, whereas the displacement of the nucleus from the cell centroid was unaffected, which shows that the defect is specific to the basal bodies (Fig. 2G, Extended Data Fig. S2B). Centrin still assembles at the basal bodies and the transition zones, and the NBBCs extend from the mispositioned basal bodies to the nucleus, so that their orientation follows that of the basal bodies. In contrast, centrin either does not localize to the DSF or appears as a stub next to the basal body (Fig. 2C,D, yellow arrowheads), consistent with previous work showing that the DSF is absent or incomplete in these mutants [32, 34]. Finally, we examined a mutant with defects at the distal end of the basal bodies that impair the transition from triplet to doublet microtubules (*uni2*) [35–37]. In *uni2* mutants, centrin is absent from the transition zone and the basal body of one cilium (Fig. 2B, yellow arrowhead), whereas the DSF, the NBBCs and the branched network surrounding the nucleus are preserved (Fig. 2B). Centrin at the transition zone and the basal body can therefore be lost without affecting the other centrin networks, including the DSF that connects the two basal bodies.

Altogether, analysis of these mutants shows that basal bodies are not required for centrin fibers to assemble, but that each basal body anchors and orients its own NBBC, and does so even in the absence of a DSF, as in *vfl3* and *asq2* cells. Centrin at the DSF behaves as a separate pool, since it localizes to the DSF whenever this structure is present, whether complete or reduced to a stub, and its loss leaves all other centrin structures intact. The same holds for centrin at the transition zone and the basal body, whose loss from one cilium in *uni2* cells leaves the DSF, the NBBCs and the branched network intact. Finally, the branched centrin network that surrounds the nucleus assembles in the absence of all other centrin structures, indicating that it forms independently of them. Having established that these centrin structures constitute separate pools, we next investigated their functions.

### CENTRIN MUTATIONS AFFECT CILIA ASSEMBLY BUT NOT BASAL BODY POSITIONING

To dissect the functions of the distinct parts of the centrin network, we turned to previously identified centrin mutants. The centrin mutant *vfl2* was first isolated on the basis of its variable number of cilia [38]. In *vfl2* cells, centrin is retained only at the basal bodies, whereas all centrin fibers are absent, including the NBBCs, the branched network surrounding the nucleus and the DSF that connects the two basal bodies [19, 27]. Strains derived from *vfl2*, in which this phenotype is partially rescued while the *vfl2* mutation is retained, were later isolated but never studied [27]. Thus, to understand the functions of the centrin networks, we first sought to characterize these strains.

We began by sequencing the centrin gene in wild type, *vfl2*, and the two derived strains (Extended Data Fig. S3A; see Methods). In *vfl2*, a G-to-A change replaces a glutamic acid with a lysine in the third EF-hand domain, as originally reported [27], although at a position that differs from the published one. The mutation was reported as E101K [27], but it corresponds to residue 102 when counting from the initiator methionine. This is a difference in numbering only, since the mutated residue is the second of two consecutive glutamic acids in both the original study and our sequence. We therefore refer to the *vfl2* mutation as E102K. Both derived strains retain E102K and carry a second point mutation in the same EF-hand domain, R97H in one and K105Q in the other (Extended Data Fig. S3A,B). We refer to them as *vfl2* +R97H and *vfl2* +K105Q, and together as the *vfl2*- derived mutants.

Next we sought to determine whether centrin localization to the different fibrillar centrin structures is restored in the *vfl2*-derived mutants. When imaged by ExM, their centrin networks are intermediate between those of wild-type and *vfl2* cells (Fig. 3A–D). In contrast with *vfl2* cells, centrin localizes to the DSF and the NBBCs in both *vfl2*-derived mutants, but these structures exhibit notable geometric defects compared with the wild type. These mutants have no defects in the positioning of the basal bodies, since the angle of the basal bodies relative to the cell-pyrenoid axis is unchanged (Fig. 3E). However, the distance between the two basal bodies is significantly larger (Fig. 3E). This suggests that cell polarity is not altered in these mutants, and that the larger distance between the basal bodies instead reflects a structural defect in the DSF, which we examine below. NBBC length is significantly reduced in *vfl2* +K105Q (median 0.58 µm) and modestly reduced in *vfl2* +R97H (median 0.70 µm) compared with wild type (median 0.78 µm), and in *vfl2* +K105Q the centrin fibers are also thinner (Fig. 3C,F). Despite this, the position of the nucleus relative to the cell centroid is comparable to that of wild-type cells in both *vfl2*-derived mutants (Extended Data Fig. S3D). These defects are not explained by a reduction in centrin levels, since both *vfl2*-derived mutants have amounts of centrin comparable to those of wild-type cells (Extended Data Fig. S3C). The *vfl2* mutant differs from the *vfl2*-derived mutants in several respects. Its basal bodies are not farther apart than in wild type, although centrin is absent from the DSF, and a subset of cells shows markedly elevated basal body angles (Fig. 3E). Its nucleus is only mildly displaced from the cell centroid, although these cells lack NBBCs altogether, suggesting that the NBBCs make at most a minor contribution to nuclear positioning (Extended Data Fig. S3D). In agreement with previous reports, centrin levels are reduced in *vfl2* cells (Extended Data Fig. S3C) [27, 31].

**FIG 3.**
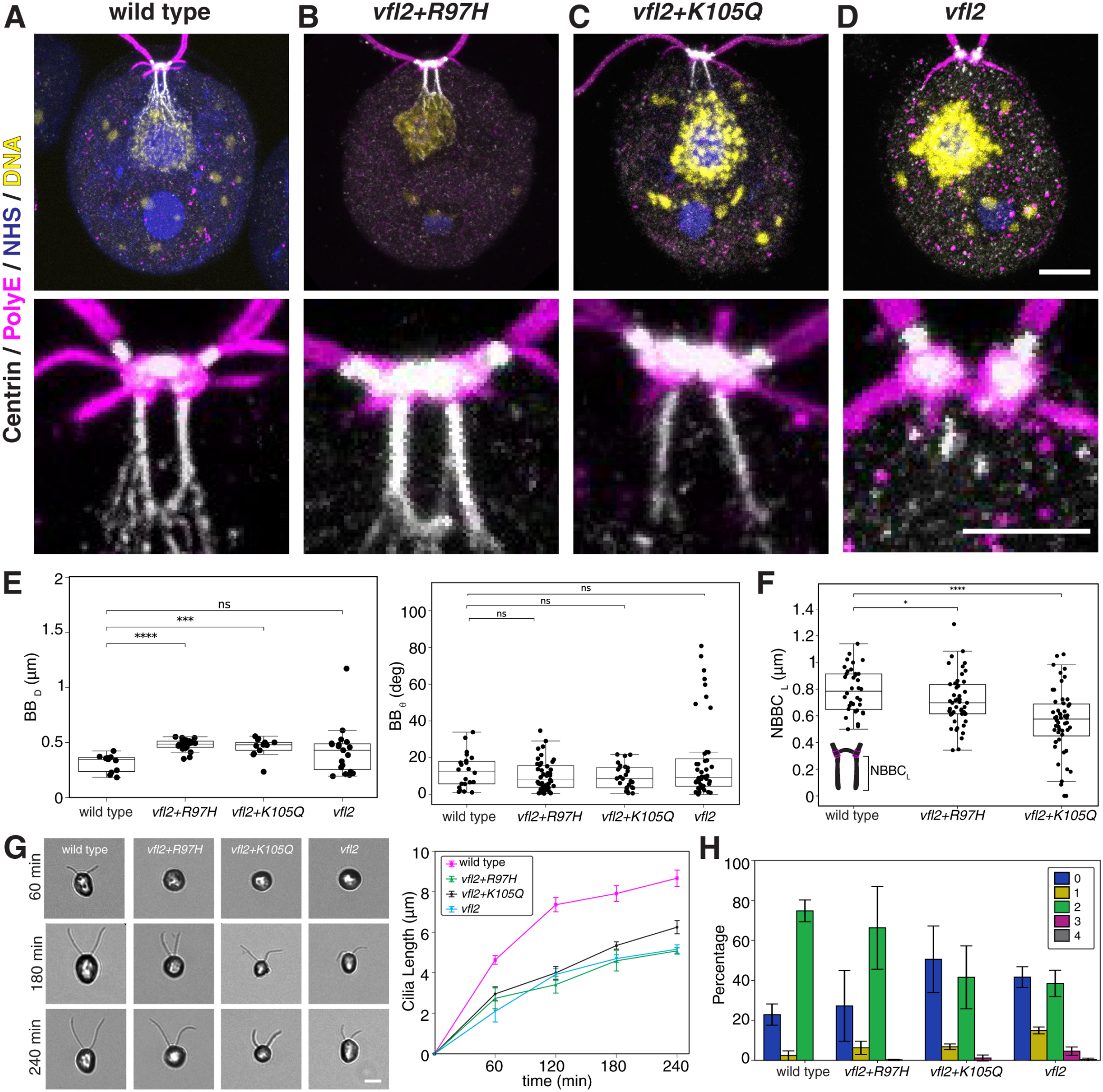
Centrin mutants show altered basal body spacing and NBBC length and defects in ciliogenesis and cilia regeneration. (A–D) Top: wild-type (A), *vfl2+R97H* (B), *vfl2+K105Q* (C) and *vfl2* (D) cells stained for centrin (white), PolyE (magenta), NHS-ester (blue) and DNA (yellow) and imaged by ExM. Bottom: inset showing the basal body region. (E) Left: distance between the basal bodies (BB_*D*_) for the four genotypes. Right: basal body angle (BB_*θ*_) for the same genotypes. Sample sizes: wild type (CC-620), *n*_cells_ = 11; *vfl2+R97H, n*_cells_ = 22; *vfl2+K105Q, n*_cells_ = 14; *vfl2, n*_cells_ = 20. (F) NBBC length (NBBC_*L*_, schematic inset) for wild-type, *vfl2+R97H* and *vfl2+K105Q* cells. Sample sizes: wild type, *n*_cells_ = 18; *vfl2+R97H, n*_cells_ = 25; *vfl2+K105Q, n*_cells_ = 29. (G) Left: representative DIC images of regenerating cilia at 60, 180 and 240 min after deciliation. Right: mean cilia length over time after deciliation. Data are mean ± s.e.m. across 4 independent experiments. Total cilia analysed across all time points: wild type, *n*_cilia_ = 3952; *vfl2+R97H, n*_cilia_ = 1786; *vfl2+K105Q, n*_cilia_ = 1689; *vfl2, n*_cilia_ = 1103. (H) Percentage of cells with 0, 1, 2, 3 or 4 cilia for each genotype. Data are mean ± s.d. across 4 independent experiments. Sample sizes: wild type, *n*_cells_ = 807; *vfl2+R97H, n*_cells_ = 828; *vfl2+K105Q, n*_cells_ = 731; *vfl2, n*_cells_ = 731. Box plots as in Fig. 2. In (E) and (F), each mutant was compared with wild type using a two-sided Mann–Whitney *U* test; \**p <* 0.05, \*\*\**p <* 0.001, \*\*\*\**p <* 0.0001; ns, not significant. Scale bars: 2 µm (A–D, top), 1 µm (A–D, bottom), 5 µm (G).

Centrins have a well-documented role in ciliogenesis and basal body segregation [10, 12, 19, 27, 38, 39]. Having characterized how centrin mutations affect the structures associated with the basal bodies, we next asked whether they also impair the assembly of cilia and alter their number. We therefore measured cilia length, regeneration kinetics, and cilia number in all three mutants. We find that all mutants have shorter cilia than wild type, with mean lengths of 7.2 and 7.1 µm in *vfl2* +R97H and *vfl2* +K105Q, respectively, and 6.3 µm in *vfl2*, compared with 9.2 µm in wild type (Extended Data Fig. S3E). Regeneration after deciliation is also slower in all mutants. At 60 min, wild-type cilia have recovered approximately half of their steady-state length, whereas cilia of the mutants have recovered only 35–40%. By 240 min, wild-type cilia have recovered their steady-state length, those of *vfl2* +K105Q and *vfl2* cells 85–90%, and those of *vfl2* +R97H cells approximately 70% (Fig. 3G, Extended Data Fig. S3E). Cilia assembly is therefore impaired in all three mutants. In contrast, cilia number distinguishes *vfl2* from the *vfl2*-derived mutants (Fig. 3H). Cells with an abnormal number of cilia, that is one, three, or four, make up approximately 20% of *vfl2* cultures but fewer than 10% of wild-type and *vfl2*-derived cultures. The second mutation therefore suppresses the variable number of cilia of *vfl2*, but not its defects in cilia assembly.

Altogether, analysis of these mutants shows that a second mutation in centrin restores the localization of centrin to the DSF and the NBBCs, although with an altered geometry. Neither this altered geometry nor the loss of the fibers in *vfl2* strongly affects the positioning of the basal bodies or the nucleus. In contrast, cilia are shorter and slower to regenerate in all three mutants, including those in which the centrin fibers are restored and centrin levels are normal, indicating that cilia assembly depends on centrin independently of these fibers. Only the variable number of cilia is specific to *vfl2*, and it could therefore result from its reduced centrin levels or from the absence of the centrin fibers. The three mutants thus share their defects in cilia assembly but differ in the centrin network at the DSF, which lacks centrin in *vfl2* and has a reduced amount in its middle in the *vfl2*-derived mutants. Because the DSF physically connects the two basal bodies, these mutants allowed us to test whether the centrin network at this fiber has a mechanical function, beyond the structural roles of centrin in cilia assembly.

### CENTRIN AT THE DISTAL STRIATED FIBER IS REQUIRED FOR CILIA COORDINATION

The DSF links the two basal bodies at their distal ends. In all mutants lacking a complete DSF described so far, the loss of the fiber is a consequence of defects in the basal bodies [32, 34, 40–42]. In *vfl3* cells, the DSF is absent and the coordination between the two cilia is disrupted [43], but because the fiber is lost entirely, the contribution of its components to coordination cannot be assessed. Addressing this question has been limited by our knowledge of the molecular composition of the DSF, of which centrin is the only known component [19, 23, 44]. The centrin mutants characterized above, in which the basal bodies are correctly positioned but centrin at the DSF is absent or altered, provide a direct test of the contribution of the fiber to cilia coordination. We therefore explored whether centrin at the DSF is required for the synchronization of cilia that underlies swimming.

We first quantified the defects of the DSF in expanded cells of the *vfl2*-derived mutants. We measured the intensity profile of centrin along the line connecting the two basal bodies and defined the width of the DSF as the length over which this profile exceeds half of its maximum (see Methods B 4). The DSF is significantly wider in both *vfl2*-derived mutants than in wild type, with a median width of approximately 0.4 µm in each mutant compared with 0.2 µm in wild type (Fig. 4A,B), in agreement with the larger distance between their basal bodies, which was measured independently from the positions of the basal bodies (Fig. 3E). The profiles also show that centrin is reduced in the middle of the fiber in both mutants, whereas it is evenly distributed along the DSF in wild type (Extended Data Fig. S4A). A similar defect in the central region of the DSF was observed by electron microscopy in other centrin mutants derived from *vfl2* [27]. In *vfl2* cells, no centrin is detected at the DSF (Fig. 4A), consistent with electron microscopy showing that these cells lack the DSF altogether [45]. Next we examined whether these defects affect swimming, which in *Chlamydomonas* depends on the coordinated, breaststroke-like beat of the two cilia [46, 47]. Wild-type cells swim along helical trajectories with a mean speed of 76.9 ± 39.4 µm/s, within the range of previous measurements (Fig. 4C,D) [46, 48]. In contrast, all centrin mutants swim significantly more slowly, with mean speeds of 38.6 ± 27.6 µm/s in *vfl2* +R97H, 43.4 ± 23.5 µm/s in *vfl2* +K105Q, and 24.8 ± 16.9 µm/s in *vfl2*, and their trajectories lack the helical signature of wild-type swimming (Fig. 4C,D). These swimming defects are consistent with a loss of cilia coordination, but they could also result from the shorter cilia of the mutants.

**FIG 4.**
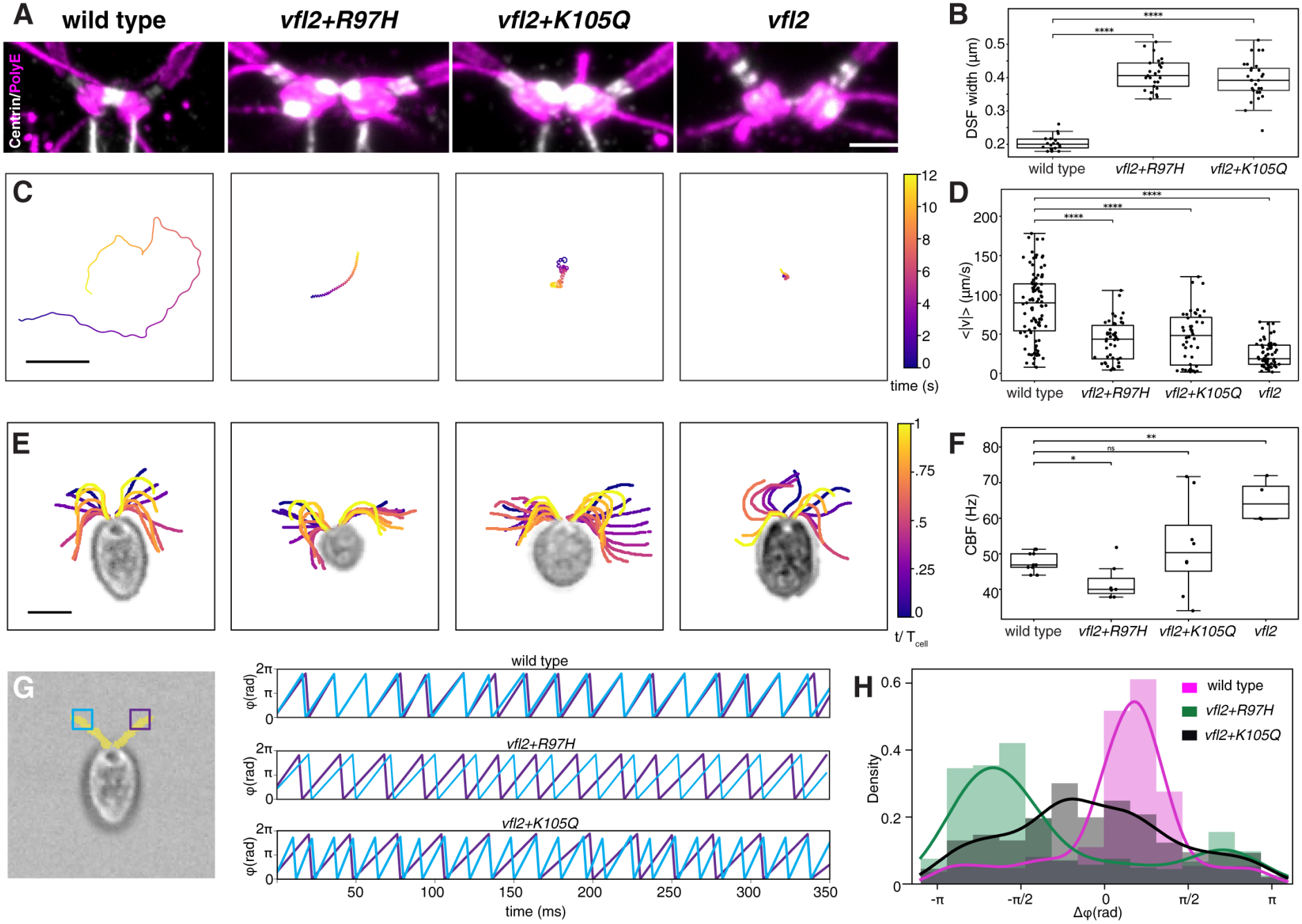
Centrin bundles at the distal striated fibers are required for cilia coordination. (A) Basal body regions of wild-type, *vfl2+R97H, vfl2+K105Q* and *vfl2* cells stained for centrin (white) and PolyE (magenta) and imaged by ExM. (B) DSF width in wild-type, *vfl2+R97H* and *vfl2+K105Q* cells. Sample sizes: wild type (CC-620), *n*_cells_ = 18; *vfl2+R97H, n*_cells_ = 25; *vfl2+K105Q, n*_cells_ = 29. (C) Representative trajectories of swimming cells for each genotype over 12 s. Color indicates time. (D) Mean swimming speed per trajectory for each genotype. Sample sizes: wild type, *n*_trajectories_ = 95; *vfl2+R97H, n*_trajectories_ = 44; *vfl2+K105Q, n*_trajectories_ = 46;*vfl2, n*_trajectories_ = 64 (E) Traced cilia of representative cells of each genotype over one beat cycle. Color indicates time normalized to the beat period (*t/T*). Time interval between consecutive traces is 2 ms. (F) Cilia beat frequency (CBF) per cilium. Only cilia that beat periodically throughout the timelapse were analysed. Sample sizes: wild type, *n*_cells_ = 5, *n*_cilia_ = 10; *vfl2+R97H, n*_cells_ = 5, *n*_cilia_ = 7; *vfl2+K105Q, n*_cells_ = 5, *n*_cilia_ = 8; *vfl2, n*_cells_ = 4, *n*_cilia_ = 4. (G) Left: DIC image of a wild-type cell overlaid with cilia traces (yellow). Squares indicate the regions of interest used to extract the ciliary signal over time. Right: phase of the two cilia over time in representative wild-type, *vfl2+R97H* and *vfl2+K105Q* cells, showing in-phase beating, phase-locking away from the in-phase state, and distinct beat frequencies, respectively. (H) Distribution of the phase difference (Δ*φ*) between the two cilia of individual cells, pooled across all time points and cells for each genotype. Bars show binned frequencies and lines show Gaussian kernel density estimates. Sample sizes: wild type, *n*_cells_ = 5; *vfl2+R97H, n*_cells_ = 2; *vfl2+K105Q, n*_cells_ = 4. Movies in (E–H) were recorded at 1000 fps and analysed over 0.5 s at 500 fps. Box plots as in Fig. 2. In (B), (D) and (F), each mutant was compared with wild type using a two-sided Mann–Whitney *U* test; \**p <* 0.05, \*\**p <* 0.01, \*\*\*\**p <* 0.0001; ns, not significant. Scale bars: 0.5 µm (A), 200 µm (C), 5 µm (E).

We therefore measured cilia coordination directly, by imaging ciliary beating in free-swimming cells with high-speed DIC microscopy and extracting the waveform, beat frequency, and phase of each cilium (Fig. 4E–H, Movie S1; see Methods). Because in-plane ciliary beating is difficult to capture in free-swimming cells, sample sizes are small (five cells each for wild type, *vfl2* +R97H, and *vfl2* +K105Q, and four cells for *vfl2*). In wild-type cells, ciliary beating is stable and symmetrical, with a mean cilia beat frequency (CBF) of 47.7 ± 2.6 Hz, and the two cilia of each cell beat at the same frequency (Fig. 4F, Extended Data Fig. S4B). Four of the five cells maintain sustained synchrony, interrupted only by short asynchronous bursts, which results in a narrow distribution of phase differences centered near zero (Fig. 4G,H). In *vfl2* +K105Q cells, the two cilia beat at markedly different frequencies, the slower at 42 ± 6 Hz and the faster at 62 ± 9 Hz, so that the mean CBF is not different from wild type despite a wide spread (Fig. 4F). The difference in frequency between the two cilia of the same cell ranges from 5 to 36 Hz, which causes a continuous phase drift and a broad distribution of phase differences (Fig. 4G,H, Extended Data Fig. S4B). In *vfl2* +R97H cells, the mean CBF is modestly reduced, to 42 ± 5 Hz (Fig. 4F). In three of the five cells, one cilium beats defectively, alternating between rigid paralysis and very slow, broad strokes, and in the two cells in which both cilia beat periodically, the cilia are phase-locked but shifted away from the in-phase state of wild-type cells (Fig. 4G,H). Finally, *vfl2* cells show the most severe phenotype. In each cell, only one cilium beats periodically, at an elevated mean CBF of 65 ± 5 Hz, and the other beats irregularly, alternating between rigid motion and broad, slow strokes that overlap with the beating region of the periodic cilium (Fig. 4E,F, Extended Data Fig. S4C).

Thus, centrin at the DSF is required for the coordination of the two cilia. Coordination is lost in all three centrin mutants, and the nature of the defect correlates with the organization of centrin at the DSF. In *vfl2*, which lacks centrin at the DSF, only one cilium beats periodically. In the *vfl2*-derived mutants, in which the DSF is wider and contains less centrin in its middle, the two cilia beat at different frequencies in *vfl2* +K105Q and are phase-locked away from the in-phase state in *vfl2* +R97H. A physical connection between the basal bodies is therefore not sufficient for cilia coordination, which instead depends on the organization of centrin within the DSF. Because centrin-based fibers contract in response to Ca^2+^, the DSF may coordinate the cilia through its mechanical properties rather than as a static link. We therefore next asked whether the DSF and the other centrin networks at the ciliary base contract in response to Ca^2+^.

### CENTRIN STRUCTURES AT THE CILIARY BASE CONTRACT IN RESPONSE TO CA^2+^

Centrin-based fibers contract in response to Ca^2+^, and in *Chlamydomonas*, the NBBCs and the transition zone have been shown to contract [26, 49, 50]. Whether the DSF also contracts, and whether the centrin mutations affect the contractility of these structures, remains unknown. To address this, we isolated nucleoflagellar apparatuses (NFAPs) from wild-type, *vfl2* +R97H, and *vfl2* +K105Q cells, incubated them without or with Ca^2+^ (5 mM free Ca^2+^), and imaged the centrin networks by ExM (Fig. 5A–C; see Methods).

**FIG 5.**
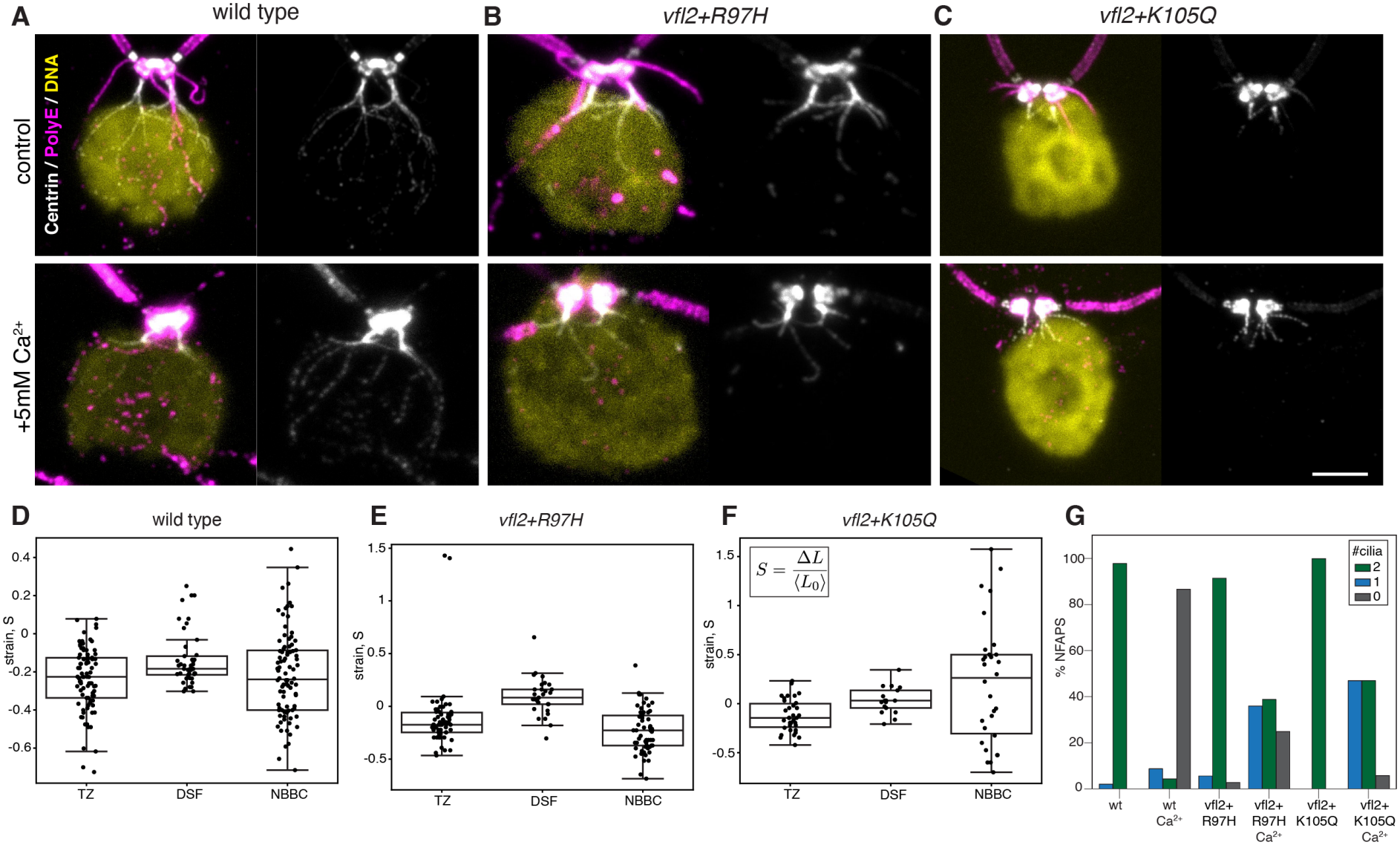
Centrin substructures contract in response to Ca^2+^, and centrin mutations differentially affect their contractility. (A) Nucleoflagellar apparatuses (NFAPs) isolated from wild-type cells, stained for centrin (white), PolyE (magenta) and DNA (yellow) and imaged by ExM, without (top) or with (bottom) Ca^2+^ treatment (5 mM free Ca^2+^). (B,C) As in (A), for *vfl2+R97H* (B) and *vfl2+K105Q* (C). (D) Ca^2+^-induced strain, *S* = Δ*L/ ⟨L*_0_*⟩*, of the transition zone (TZ), distal striated fiber (DSF) and nucleus–basal body connectors (NBBCs) in wild-type NFAPs. *⟨L*_0_*⟩* is the mean length of each structure in untreated NFAPs of the same genotype. Negative values indicate contraction. (E,F) As in (D), for *vfl2+R97H* (E) and *vfl2+K105Q* (F). (G) Percentage of NFAPs retaining 0, 1 or 2 cilia without and with Ca^2+^ treatment for each genotype. Sample sizes (D–F): wild type (CC-620), *n*_NFAPs_ = 45; *vfl2+R97H, n*_NFAPs_ = 29; *vfl2+K105Q, n*_NFAPs_ = 17. (G) wild type control, *n*_NFAPs_ = 47; wild type calcium, *n*_NFAPs_ = 45;*vfl2+R97H* control, *n*_NFAPs_ = 35; *vfl2+R97H* calcium, *n*_NFAPs_ = 36;*vfl2+K105Q* control, *n*_NFAPs_ = 19; *vfl2+K105Q* calcium, *n*_NFAPs_ = 17;Box plots as in Fig. 2. Scale bar: 1 µm (A-C).

In wild-type NFAPs, the extended centrin network collapses into a compact cluster around the basal bodies after Ca^2+^ treatment (Fig. 5A). To quantify the contraction of each structure, we measured its Ca^2+^-induced strain, *S* = Δ*L/ ⟨L*_0_*⟩*, where *⟨L*_0_*⟩* is the mean length of the structure in untreated NFAPs of the same genotype (Fig. 5D–F; see Methods). In wild type, the strain is negative at the TZ, the DSF, and the NBBCs, with medians of approximately −0.2 for all three structures, showing that each of them contracts by about 20% in response to Ca^2+^ (Fig. 5D). In both *vfl2*-derived mutants, the TZ still contracts, whereas the DSF does not, with a strain distribution shifted toward positive values in *vfl2* +R97H and centered near zero in *vfl2* +K105Q (Fig. 5E,F). The two mutations differ in their effect on the NBBCs. In *vfl2* +R97H, the NBBCs contract to a similar extent as in wild type, whereas in *vfl2* +K105Q, in which they are also shorter and thinner (Fig. 3F), they show a broad and often positive strain distribution, indicating impaired contraction (Fig. 5E,F). Ca^2+^ treatment does not reduce the size of the nucleus in any of the three genotypes, and the nucleus even enlarges in *vfl2* +R97H NFAPs (Extended Data Fig. S5), indicating that the Ca^2+^-induced contraction of the centrin networks does not compress the nucleus.

Ca^2+^ treatment also induces cilia shedding, a process that is accompanied by contraction of the centrin-containing fibers of the TZ [26, 50]. We therefore asked whether shedding is affected in the *vfl2*-derived mutants, in which the TZ still contracts (Fig. 5E,F). After Ca^2+^ treatment, 87% of wild-type NFAPs shed both cilia and 9% shed one (Fig. 5G). In contrast, shedding is incomplete in both *vfl2*-derived mutants. Only 25% of *vfl2* +R97H NFAPs shed both cilia and 36% shed one, and only 6% of *vfl2* +K105Q NFAPs shed both cilia and 47% shed one (Fig. 5G). Contraction of the TZ is therefore not sufficient for Ca^2+^-induced cilia shedding, which also depends on centrin in other structures or on centrin functions other than contraction.

Together, these measurements show that the TZ, the DSF, and the NBBCs each contract in response to Ca^2+^, and that the centrin mutations affect the contractility of each structure differently. DSF contraction is lost in both *vfl2*-derived mutants, the same mutants in which the DSF is wider and the two cilia are not coordinated (Fig. 4), consistent with a mechanical role of centrin at the DSF in cilia coordination.

## DISCUSSION

Here, we show that centrin builds structurally distinct substructures at the *Chlamydomonas* basal body apparatus, and that these substructures differ in how they assemble and which ciliary functions they support. The NBBCs and the branched network surrounding the nucleus assemble even in the absence of basal bodies, but they rely on basal bodies to be anchored and oriented within the cell, whereas centrin at the DSF is lost when the basal bodies are mispositioned. Point mutations in centrin reveal how the integrity of these structures relates to their function. In *vfl2* cells, in which centrin levels are reduced, all centrin fibers are lost, including the branched network surrounding the nucleus, whereas the *vfl2*-derived mutants still assemble NBBCs and a DSF, albeit with altered geometry. Despite these structural differences, all three mutants have defects in cilia assembly and regeneration, suggesting that this function does not depend on the centrin fibers. Contractility, by contrast, is sensitive to the organization of the fibers. In wild-type cells, the TZ, the NBBCs and the DSF all contract by approximately 20% in response to Ca^2+^. In both *vfl2*-derived mutants, the TZ still contracts, whereas the DSF, which is wider, no longer contracts, and the NBBCs lose their contractility in *vfl2* +K105Q, in which they are also shorter and thinner (Fig. 3F), but not in *vfl2* +R97H. The loss of a contractile DSF coincides with the loss of coordination between the two cilia. Altogether, these results show that centrin is a building block whose assembly into filamentous structures is required for their contractility and, at the DSF, for the coordination of the two cilia.

In unicellular eukaryotes, centrin carries out a wider range of functions than in metazoans, and in ciliates these functions are divided among distinct centrin isoforms. In *Tetrahymena*, only one of four centrins, Cen1, is essential for basal body duplication and maintenance [9]. In *Paramecium*, basal body-associated centrins control the geometry of basal body duplication [8], whereas a distinct subfamily of centrins assembles the contractile infraciliary lattice, which positions and anchors the basal bodies [51, 52]. In *Chlamydomonas*, Vfl2 centrin participates in basal body duplication and segregation [7, 19, 27, 38], and we find that it is also required for the contraction of the DSF and for its function in cilia coordination. Our results suggest that a single protein can fulfill these diverse roles. Rather than being divided among isoforms, the functions of Vfl2 centrin are divided among structures, with the same protein forming compact accumulations at the basal bodies and TZs, linear fibers such as the NBBCs and the DSF, and a branched network surrounding the nucleus.

We find that each of these structures assembles independently. Basal bodies are not required for centrin to assemble into fibers. In *bld10* mutants, which lack basal bodies, centrin still forms NBBC-like fibers and a branched network surrounding the nucleus, in agreement with the earlier observation that NBBCs form in *bald-2* cells, which retain only rudimentary basal bodies [31]. Basal bodies instead determine where these fibers are placed. In *bld12* cells, NBBC-like filaments are correctly positioned only when a basal body remnant is present, and in *vfl3* and *asq2* mutants, in which basal bodies assemble but are mispositioned, centrin at the DSF is absent or reduced to a stub. Network assembly therefore appears to proceed in two tiers, in which centrin fibers form autonomously and are anchored and oriented by the basal bodies, whereas centrin at the DSF depends on correctly positioned basal bodies.

Beyond organizing the basal body apparatus, centrin contributes to the assembly of the cilia it templates. The Vfl2 centrin mutants more often lack cilia, assemble shorter cilia and regenerate them more slowly, mirroring the requirement for centrins in primary ciliogenesis in human cells [10, 12, 39]. In human cells, this requirement depends on Ca^2+^ binding, as mutation of any single EF-hand domain of centrin 2 impairs ciliogenesis [39], and the mutations studied here likewise lie in an EF-hand domain. Which centrin population underlies these defects remains unclear. Besides its localization at the basal bodies and the TZ, centrin is present in the axoneme as a light chain of a subset of inner-arm dyneins [53], and the point mutations affect all three populations at once. Distinguishing their contributions will require perturbing centrin locally, for example by targeting its binding partners at each of these substructures.

Cilia coordination, in contrast, can be linked more directly to a single substructure, the DSF. Because hydrodynamic coupling between the two cilia is too weak to account for their synchrony [54], and because mutants lacking the fibers that connect the basal bodies lose synchrony [43], the cilia are thought to be coupled through the basal bodies. Our results show that an organized centrin bundle at the DSF is required for cilia coordination. In *vfl2* +R97H and *vfl2* +K105Q cells, centrin localizes at the DSF in an abnormal pattern and the two cilia fail to coordinate. A connection between the basal bodies therefore appears not to be sufficient, and coupling instead depends on the organization of the centrin bundle at the DSF. When uncoupled, the *cis* and *trans* cilia beat at different intrinsic frequencies [55, 56]. In *vfl2* +K105Q cells, the two cilia beat at approximately 42 and 62 Hz, a difference comparable to that between these intrinsic frequencies, suggesting that each cilium beats at its own frequency because the coupling between them is lost, rather than because the cilia themselves are defective. Because centrin is also associated with inner-arm dyneins [53], however, a contribution of axonemal centrin cannot be excluded. Measuring coupling strength directly in the centrin mutants, for example by entraining the cilia of micropipette-held cells with external flows [54, 56], will establish how the structure of the DSF contributes to coupling.

Cilia coordination, in contrast, can be linked more directly to a single substructure, the DSF. Because hydrodynamic coupling between the two cilia is too weak to account for their synchrony [54], and because mutants lacking the fibers that connect the basal bodies lose synchrony [43], the cilia are thought to be coupled through the basal bodies. Our results show that this coupling requires an organized centrin bundle at the DSF. In *vfl2* +R97H and *vfl2* +K105Q cells, centrin localizes to the DSF in an abnormal pattern, and the two cilia fail to coordinate, indicating that a physical connection between the basal bodies is not sufficient for coordination. When uncoupled, the *cis* and *trans* cilia beat at different intrinsic frequencies [55, 56]. In *vfl2* +K105Q cells, the two cilia beat at approximately 42 and 62 Hz, a difference comparable to that between these intrinsic frequencies, suggesting that each cilium beats at its own frequency because the coupling between them is lost, rather than because the cilia themselves are defective. However, since centrin is also associated with inner-arm dyneins [53], a contribution of axonemal centrin cannot be excluded. Measuring coupling strength directly in the centrin mutants, for example by entraining the cilia of micropipette-held cells with external flows [54, 56], will establish how the structure of the DSF contributes to coupling.

The organization of the DSF may matter because it determines how the fiber responds to Ca^2+^. Centrin-based fibers contract in response to Ca^2+^ across green algae, including the striated root fibers of *Tetraselmis*, the distal connecting fiber of *Spermatozopsis*, and the NBBCs and TZs of *Chlamydomonas* [18, 19, 25, 26, 49]. We find that the DSF of *Chlamydomonas* contracts as well, and that contraction is lost where the structure of the fibers is altered. In both *vfl2*-derived mutants, the TZ still contracts, whereas the wider DSF does not, and the NBBCs contract in *vfl2* +R97H but not in *vfl2* +K105Q, in which they are shorter and thinner. Because the same mutant protein contracts in one structure but not in another, this defect cannot reflect a general loss of Ca^2+^ binding, and instead suggests that contraction depends on the organization of centrin within each fiber. At the DSF, the loss of contractility coincides with the loss of coordination, suggesting that the mechanical state of the DSF, and not only its presence, contributes to coupling between the two cilia. By controlling the mechanical state of the DSF, Ca^2+^ signaling could thereby tune ciliary synchrony.

Our shedding measurements also address the long-debated role of centrin in cilia shedding. Previous studies showed that Ca^2+^-induced contraction of the centrin-containing stellate fibers of the TZ accompanies shedding, and that anti-centrin antibodies block shedding in detergent-extracted cell models [26, 50]. Yet *vfl2* cells were found to shed their cilia normally [45, 57, 58]. We find that the outcome depends on the assay. In isolated NFAPs treated with Ca^2+^, the *vfl2*-derived mutants shed their cilia incompletely, even though their TZs still contract, whereas living cells of all three mutants deciliate completely after pH shock (Fig. 3G). In isolated NFAPs, incomplete shedding is therefore associated with mutant centrin, and TZ contraction alone is not sufficient for shedding. Whether centrin acts directly in shedding, or whether the mutations impair shedding indirectly, for example by altering the structure of other centrin networks, remains unclear. The difference between NFAPs and living cells may reflect components of the deciliation machinery that are active only in living cells, but the two assays also use different stimuli, and comparing them under identical conditions will be needed to resolve this.

How Vfl2 centrin assembles into substructures with such diverse architectures, from compact bundles to branched networks, remains the central open question. Candidate determinants are centrin-binding partners specific to each substructure. Sfi1 binds multiple centrin molecules at the yeast spindle pole body [14] and at the distal end of human centrioles [10], and Sfi1-like proteins build the contractile infraciliary lattice of *Paramecium* [15]. Sfi1 homologs have been identified in the genome of the green alga *Ostreococcus* [59], but whether *Chlamydomonas* Sfi1-like proteins contribute to the DSF or the NBBCs remains to be determined. Identifying these partners could explain how centrin forms structures with distinct architectures, and how mutations in centrin alter their organization. A second possibility is that additional centrin-like proteins contribute to some of these structures. Although *Chlamydomonas* centrin is encoded by a single-copy gene [60], the *vfl2* phenotype led to the proposal that *Chlamydomonas* contains an additional centrin gene implicated in basal body duplication [61]. Whether such a protein exists, and whether it contributes to the residual centrin signal at the basal bodies of *vfl2* cells, remains to be determined.

More broadly, our results show that a single, conserved Ca^2+^-binding protein can be assembled into structurally and functionally distinct networks within one cell. Much like actin and tubulin, whose functions are defined by the architectures they form, centrin acquires its functions through the assemblies it builds, establishing it as a cytoskeletal element in its own right. At the ciliary base, the DSF, acts as a contractile coupler between the two cilia, whose function depends on how centrin is organized within it.

## METHODS

### A. Cell preparation and imaging

#### 1. Strains and culture conditions

All strains were obtained from the Chlamydomonas Resource Center (https://www.chlamycollection.org/). The centrin mutant *vfl2* (CC-2530) and the *vfl2*-derived mutants *vfl2* +R97H (CC-3015) and *vfl2* +K105Q (CC-3014) [27] were compared with the wild-type strain CC-620. The basal body mutants *vfl3* (CC-1686) and *asq2* (CC-4301) were compared with the wild-type strain CC-124. The basal body mutants *bld10, uni2* and *bld12* correspond to strains CC-4076, CC-4183 and CC-4162 respectively. Cells were maintained on 1.5% TAP agar plates at 20 ^*°*^C under a 14 h/10 h light/dark cycle. For experiments, cells were inoculated into liquid TAP or M medium, as indicated below, and grown overnight under constant light with shaking at 150 rpm.

#### 2. Immunofluorescence

Cells were washed, resuspended in PBS and allowed to adhere for 3 min to coverslips coated with 0.1% poly-D-lysine. Adhered cells were fixed for 20 min in 1% paraformaldehyde (PFA) and 0.1% Triton X-100 in PHEM buffer (60 mM PIPES, 25 mM HEPES, 10 mM EGTA, 2 mM MgCl_2_, pH 7.0), and further permeabilized twice for 5 min in methanol at −20 ^*°*^C. After three 5-min washes in PBS, samples were blocked for 30 min in 5% BSA, 5% goat serum and 1% cold-water fish gelatin in PBS, and incubated overnight at 4 ^*°*^C with rabbit anti-polyglutamate chain (PolyE, 1:2000; AdipoGen AG-25B-0030-C050) and mouse anti-centrin (1:400; Milli-pore ZMS1054) diluted in blocking buffer. Samples were washed three times for 5 min in PBS with 0.1% Triton X-100 and incubated for 1 h at room temperature in the dark with Alexa Fluor 568 goat anti-rabbit and Alexa Fluor 488 goat anti-mouse (Invitrogen A11001 and A11004) secondary antibodies (1:800 in blocking buffer). After two washes in PBS with 0.1% Triton X-100 and one wash in PBS, samples were mounted in VectaShield HardSet Antifade Mounting Medium with DAPI (Vector Laboratories H-1500-10).

#### 3. Ultrastructure expansion microscopy (U-ExM)

U-ExM was performed as described [62], with minor modifications. Cells grown overnight in TAP under constant light were pelleted at 800 × *g* for 5 min, washed and resuspended in PBS, and crosslinked for 5 h at 37 ^*°*^C in 2% acrylamide and 1.4% formaldehyde in PBS. A small drop of concentrated cells was placed on a 12-mm round coverslip and gelled at 4 ^*°*^C in monomer solution (19% (wt/wt) sodium acrylate, Sigma-Aldrich 408220; 10% (wt/wt) acrylamide, Sigma-Aldrich A4058; 0.1% (wt/wt) N,N’-methylenebisacrylamide, Sigma-Aldrich M1533 in PBS) supplemented immediately before use with 0.5% (wt/wt) APS (ThermoFisher 17874) and 0.5% (wt/wt) TEMED (ThermoFisher 17919). Gels were polymerized for 1 h at 37 ^*°*^C in a humid chamber, detached from the coverslips and denatured for 2 h at 95 ^*°*^C in denaturation buffer (50 mM Tris, 200 mM NaCl, 200 mM SDS, pH 9.0). Gels were expanded overnight at 4 ^*°*^C in Milli-Q water and inspected by 20× DIC imaging to confirm expansion and identify regions of high cell density. These regions were excised, shrunk by two 15-min washes in PBS and incubated overnight at 37 ^*°*^C with shaking with anti-PolyE (1:400) and anti-centrin (1:400) in antibody solution (3% BSA, 0.1% Triton X-100 in PBS). Gels were washed three times for 10 min in PBS with 0.1% Triton X-100 and incubated for at least 5 h at 37 ^*°*^C with Alexa Fluor 488 goat anti-mouse (Invitrogen A11001) and Alexa Fluor 568 anti-rabbit secondary antibodies (1:1000 in antibody solution). For NHS-ester staining, gels were washed in 0.1 M sodium bicarbonate (pH 8.3) and incubated for 1.5 h at room temperature in the dark with NHS-ester 405 (1:500 in 0.1 M sodium bicarbonate). DNA was stained for 10 min with NIR694 nuclear dye (1:700; Biotracker SCT118). Gels were washed in PBS for 5 min, fully re-expanded by three 15-min incubations in Milli-Q water, and mounted on APTES ((3-aminopropyl)-triethoxysilane)-coated coverslips in a custom 3-mm-high acrylic chamber filled with Milli-Q water to prevent shrinking.

#### 4. Isolation and Ca^2+^ treatment of nucleoflagellar apparatuses

Nucleoflagellar apparatuses (NFAPs) were isolated from 10 ml of cells grown overnight in M medium. Cells were harvested at 1500 × *g* for 5 min. Walled strains were treated with autolysin for 30 min to remove the cell wall. Cells were resuspended in 1 ml nucleus buffer (NB; 6.7 mM Tris-HCl pH 7.3, 3.7 mM EGTA, 2.5 mM KCl, 10 mM MgCl_2_), layered on 10 ml of 30% Percoll in NB and pelleted at 500 × *g* for 5 min. The pellet was resuspended in 200 µl NB, and 100 µl was allowed to adhere for 5 min to a poly-lysine-coated coverslip. Cells were lysed for 2 min with shaking in 1% IGEPAL in NB to release the NFAPs, washed once in NB and incubated for 5 min in NB alone (control) or in NB supplemented with 9 mM CaCl_2_, giving 5 mM free Ca^2+^ in the presence of 3.7 mM EGTA. NFAPs were then crosslinked in 2% acrylamide and 1.4% formaldehyde in PBS for 5 h at 37 ^*°*^C and processed for U-ExM as described above.

#### 5. Confocal imaging

Non-expanded samples were imaged on a Nikon Ti2 inverted microscope equipped with an AX point-scanning confocal, using a Plan Apo 60 × NA 1.42 oil-immersion objective and the galvano scanner. Expanded samples were imaged on the same microscope with a 40× Apo LWD NA 1.15 water-immersion objective (pixel size 0.103 µm). High-resolution images of selected structures, including basal bodies used to determine the expansion factor, were acquired with the NSPARC detector in super-resolution mode (pixel size 0.024 µm). Z-stacks were converted to maximum-intensity projections for analysis.

#### 6. Immunoblotting

Cells from 10 ml cultures were pelleted at 800 ×*g* for 5 min, washed once in PBS and lysed in ten pellet volumes of lysis buffer (40 mM Tris pH 7.5, 5 mM MgCl_2_, 2% SDS, 1 mM PMSF and protease inhibitor cocktail). Lysates were mixed with 5× Laemmli buffer, heated for 8 min at 85 ^*°*^C and cleared by centrifugation at 10,000 rpm for 10 min. Proteins were separated by SDSPAGE and wet-transferred to nitrocellulose for 1 h at 4 ^*°*^C (25 mM Tris, 192 mM glycine, 20% ethanol). Total protein was visualized with Ponceau S. Membranes were blocked for 1 h in 3% BSA in TBST (20 mM Tris pH 7.6, 150 mM NaCl, 0.1% Tween-20) and incubated overnight at 4 ^*°*^C with mouse anti-centrin (1:5000; Milli-pore ZMS1054) and rabbit anti-AtpB (1:5000; Agrisera AS05 085) as loading control. After two 15-min washes in TBST, membranes were incubated for 1 h at room temperature with HRP-conjugated anti-mouse or anti-rabbit secondary antibodies (1:10,000), washed twice for 15 min and developed by chemiluminescence.

#### 7. PCR and Sanger sequencing

Genomic DNA was isolated from *Chlamydomonas* using QuickExtract DNA extraction solution (LGC, QE09050) as per the manufacturers’ instruction. The primer pair 5’-ATCAAGAAGATGATCTCGGAG-3’ and 5’-TCACTCACCGGATGAACTCG-3’ was used to amplify a 550bp long region surrounding the positions of the VFL2 mutation from the genomic DNA. Amplification was carried out as follows: initial denaturation at 98^*°*^C for 2 min, followed by 35 cycles of denaturation at 98^*°*^C for 30 seconds, 60^*°*^C for 30 seconds, and 72^*°*^C for 30 seconds with a final extension of 10 min. The PCR products were purified using Macherey-Nagel NucleoSpin Gel and PCR Clean-up Kit (740609.250). Sanger Sequencing of these products was performed at Microsynth AG, Switzerland.

#### 8. Cilia regeneration assay

Cells from 20 ml cultures grown overnight in M medium were pelleted at 800 ×*g* for 5 min and resuspended in 5 ml fresh TAP. Before deciliation, an aliquot was fixed in 1% glutaraldehyde to measure steady-state cilia length and number. Cells were deciliated by pH shock, adding 390 µl of 0.5 N acetic acid for 45 s followed by 420 µl of 0.5 N KOH. Cells were pelleted at 800 ×*g* for 5 min, resuspended in 5 ml fresh TAP and allowed to regenerate their cilia under light. Aliquots were collected at 60, 120, 180 and 240 min after deciliation and fixed overnight at 4 ^*°*^C in 1% glutaraldehyde in TAP. Fixed cells were allowed to adhere to poly-D-lysine-coated coverslips and imaged by DIC on a Nikon Ti2 inverted microscope with a 40× Apo LWD NA 1.15 water-immersion objective.

#### 9. Live imaging of swimming cells and ciliary beating

Cells were pelleted at 800 × *g* for 5 min and resuspended in fresh TAP at a density suitable for single-cell imaging. Imaging chambers were assembled from an uncoated glass slide and an acrylamide-coated coverslip separated by double-sided tape (40 µm spacer). To record swimming trajectories, cells were imaged by DIC on an inverted Nikon Ti2 widefield microscope with a 20× Plan Apo NA 0.8 air objective at 83 fps for 30 s. To record ciliary beating, cells were imaged in the same chambers by DIC with a 40× Apo LWD NA 1.15 water-immersion objective and a Phantom T2540 high-speed camera at 1000 fps for 2 s.

### B. Image analysis

Image analysis was performed with custom Python scripts using NumPy, SciPy, scikit-image, pandas and trackpy [63], unless stated otherwise. All segmentation masks were inspected and, where necessary, manually corrected in Fiji.

#### 1. Expansion factor

To convert measurements in expanded samples to preexpansion dimensions, the expansion factor of each experiment was calculated from the diameter of basal bodies, which have a known diameter of 200 nm [4]. Basal bodies imaged in top view with the NSPARC detector were cropped, background-subtracted using a Gaussian-filtered copy of each crop (*σ* = 20 px), smoothed (Gaussian, *σ* = 1.5 px) and normalized between their 1st and 99th intensity percentiles. The center of each basal body ring was first estimated from the centroid of the pixels above the 95th intensity percentile. Radial intensity profiles were then sampled from this center at 360 angles, between 5 and 35 px, and smoothed along the radial direction (Gaussian, *σ* = 2 samples). At each angle, the radius of maximum intensity was taken as the position of the basal body wall, and a circle was fitted to these positions by robust least squares. The expansion factor was calculated as the mean fitted diameter divided by 200 nm, and was used to convert all ExM measurements of the same experiment to pre-expansion dimensions. For mutants that lack basal bodies (*bld10* and *bld12*) and for *Tetraselmis*, the expansion factor was instead calculated from the size of the gel, as the diameter of the expanded gel divided by the measured diameter of the gel before expansion.

#### 2. Segmentation

##### Nuclei

Nuclei were segmented from the DNA channel by automatic instance segmentation with micro-SAM (model `vit_b_lm`) [64], retaining objects of 4000– 20,000 px. When micro-SAM returned no object, the nucleus was instead segmented by Otsu thresholding of the median-filtered image (10 px), followed by size filtering and hole filling.

##### Cell body and pyrenoid

Because NHS-ester labels total protein, it was used to define both the outline of the cell and the pyrenoid, which has the highest protein density. In whole cells, the cell body was segmented from the NHS-ester channel at 0.6 times the Otsu threshold, retaining objects of 10^4^–5 × 10^5^ px, followed by hole filling and median smoothing (10 px). The pyrenoid was segmented from the same channel at twice the Otsu threshold, retaining objects of 300–50,000 px with an eccentricity of at most 0.6.

##### Basal bodies and centrin structures in NFAPs

In NFAPs, basal bodies were segmented from the PolyE channel with micro-SAM, retaining objects with an aspect ratio of at most 2 and an area of 500–10,000 px. Centrin structures were segmented from the median-filtered centrin channel (3 px) at 1.5 times the Otsu threshold, retaining objects of 400–10,000 px, followed by hole filling.

#### 3. Cell geometry

To quantify the position of the basal bodies and of the nucleus, all measurements were made relative to a cell axis defined by the line connecting the centroid of the cell body to the centroid of the pyrenoid.

##### Basal body localization

In expanded whole cells, basal bodies were localized in the PolyE channel. To restrict detection to the basal body region, the channel was median-filtered (5 px) and limited to the centrin-positive region, defined by the Otsu threshold of the medianfiltered centrin channel (10 px). The restricted image was smoothed (Gaussian, *σ* = 1 px), and basal bodies were detected with the trackpy locate function (diameter 3 px, up to three features per cell). All detections were verified visually.

##### Basal body angle and distance

The angle of each basal body was calculated as the angle between the cell axis and the line connecting the cell-body centroid to the basal body, folded into the range 0–90°. The distance between basal bodies was calculated as the Euclidean distance between their positions. In cells with more than two basal bodies, the shortest pairwise distance was used.

##### Nuclear position

Nuclear displacement was calculated as the distance between the centroids of the nucleus and of the cell body. The distance between the nucleus and the basal bodies was calculated as the distance between the nuclear centroid and the mean position of the basal bodies. All distances and areas were converted to pre-expansion values using the pixel size and the expansion factor of each experiment.

#### 4. Centrin substructure measurements

The dimensions of the TZs, DSFs and NBBCs were measured from centrin intensity profiles along each structure, in expanded cells and NFAPs. Before profiles were extracted, the centrin channel was background-subtracted using a Gaussian-filtered copy (*σ* = 20 px), smoothed (Gaussian, *σ* = 2 px) and normalized between its 1st and 99th intensity percentiles. Centrin objects overlapping the basal body mask were assigned to the NBBCs, and the remaining centrin objects to the TZs. All lengths were converted to pre-expansion values using the pixel size and the expansion factor of each experiment.

##### NBBCs

To follow the curved path of each NBBC, its mask was skeletonized, and the longest path between skeleton endpoints was fitted with a smoothing spline. The portion of the spline inside the basal body mask was removed, and the spline was oriented to start at the basal body. Centrin intensity was sampled along the spline and averaged across a strip ±10 px wide (21 samples perpendicular to the spline). The resulting profile was smoothed with a Savitzky–Golay filter (window 21, first order), and NBBC length was defined as the extent of the profile above 0.6 of the normalized intensity.

##### TZs

For each basal body and its TZ, a 15-px-wide line profile was taken along the direction from the basal body centroid to the TZ centroid, starting at the edge of the basal body mask and extending 80 px beyond the TZ centroid. Profiles were smoothed with a Savitzky–Golay filter (window 11, first order), and TZ length was defined as the extent of the profile above 0.5 of the normalized intensity.

##### DSFs

DSF width was measured along the axis connecting the centroids of the two basal bodies. A 10-pxwide line profile parallel to this axis, extending 30 px beyond each basal body centroid, was placed at the perpendicular offset that maximized its mean intensity, so that the profile ran along the DSF. This offset was found by a coarse search ( ±50 px in 10-px steps) followed by a fine search ( ±5 px in 1-px steps) around the coarse optimum. The profile was normalized to the maximum of the background-subtracted image, and DSF width was defined as the extent of the profile above 0.5. The same profiles were used to compare the distribution of centrin along the DSF between genotypes (Extended Data Fig. S4A).

#### 5. Ca^2+^-induced strain

To compare the contraction of structures of different sizes, Ca^2+^-induced contraction was quantified as the strain *S* = Δ*L/ ⟨L*_0_*⟩* = (*L* − *⟨L*_0_*⟩*)*/ ⟨L*_0_*⟩*, where *L* is the length of a structure in a Ca^2+^-treated NFAP and *⟨L*_0_*⟩* is the mean length of the same structure in untreated NFAPs of the same genotype. Strain was calculated for each DSF and for each individual TZ and NBBC. Negative values indicate contraction and positive values indicate extension.

#### 6. Nuclear shape in NFAPs

To compare nuclear shape between conditions, nuclear masks from NFAPs were translated so that their centroids coincided. Nuclear area was calculated from each mask, and nuclear diameter from a least-squares circle fit to the mask contour. The average nuclear shape for each condition was obtained as the zero level set of the mean signed distance field of all aligned masks.

#### 7. Cilia number and length

Cells were identified in DIC images using probability maps generated in ilastik (v1.4.1) [65]. Only isolated cells fully contained within the field of view were analysed; clustered, truncated or ambiguous cells were excluded. The number of cilia per cell was annotated manually on the original images overlaid with the segmentation masks. Cilia length was measured with a semi-automated Python pipeline in which segmented cilia were refined, skeletonized and reduced to single centerlines. Length was calculated as the arc length of the centerline, with a local correction near the ciliary base. All measurements were reviewed manually, and failed or ambiguous segmentations were excluded or re-measured by manual tracing. For cilia number, the proportion of cells with 0–4 cilia was calculated for each biological replicate and reported as the mean ± s.d. across replicates. For regeneration, the mean cilia length at each time point was calculated for each of four biological replicates and reported as the mean ± s.e.m. across replicates.

#### 8. Ca^2+^-induced cilia shedding

The number of cilia retained by each NFAP (0, 1 or 2) was scored by visual inspection of the ExM images of control and Ca^2+^-treated samples. A cilium was scored as shed when its axoneme was completely detached from the basal body, lying at a distance from it and at an angle to the basal body axis, and as retained when the axoneme remained continuous with the transition zone. The percentage of NFAPs in each class was then calculated for each genotype and condition. The number of NFAPs analyzed per genotype and condition is given in the legend of Fig. 5.

#### 9. Cell tracking and swimming speed

To isolate moving cells from the static background, the mean of all frames of each movie was subtracted from each frame, and the absolute value of the result was smoothed with a Gaussian filter (*σ* = 1 px). Cells were then localized in each frame with trackpy [63], using a feature diameter of 15 px and a minimum integrated intensity of 20,000, and linked into trajectories with a maximum displacement of 10 px between frames and a memory of 2 frames. To retain only sustained swimming, trajectories shorter than 1000 frames (12 s) were discarded, as were trajectories with an end-to-end displacement below 10 px, which correspond to immotile cells, or above 2000 px, which result from linking errors. The remaining trajectories were inspected by overlay on the movies, and erroneous trajectories were removed manually. Gaps in the trajectories were filled by linear interpolation, and the trajectories were smoothed with a Savitzky–Golay filter (window 15 frames, second order). Instantaneous speed was calculated from frame-to-frame displacements, using a pixel size of 1.294 µm and a frame interval of 12 ms, and the mean speed of each trajectory was used for comparisons between strains.

#### 10. Ciliary beat frequency and phase

To correct for cell motion, binary masks of the cell body were generated by Gaussian blurring (*σ* = 20 px) followed by IsoData thresholding, and image sequences were registered using the spam.DIC library [66]. Incremental transformations were computed between frames separated by 5–10 ms, combined cumulatively and interpolated to yield a smooth, continuous transformation. Registered sequences were downsampled to 500 fps and rotated to align the long axis of the cell vertically. Cilia were traced manually over 0.5 s per cell. Frames with a low signal-to-noise ratio or partial out-of-plane ciliary motion, in which the ciliary position was ambiguous, were not traced. Traces were converted into binary masks after a minor dilation in Fiji.

Ciliary beat frequencies (CBFs) were quantified in Python. After the vertical alignment of the cell was refined, two symmetric regions of interest (ROIs, 2.3 × 2.3 µm) were placed 7 µm apart, one on each side of the cell. Their distance from the cell body was adjusted so that each cilium crossed its ROI only at the end of the power stroke and not during the recovery stroke. The standard deviation of the ciliary mask pixel intensities within each ROI was recorded over time, and the CBF was defined as the dominant peak of the Fourier transform of the mean-subtracted signal. Measurement uncertainty was estimated as half the full width at half maximum of the dominant peak in the normalized power spectrum. For wild-type cells, CBFs were measured during in-phase beating and excluded short transient transitions.

The phase difference between the two cilia of a cell was calculated following [43]. Motion signals were smoothed with a moving-average filter, and signal peaks were identified. The instantaneous phase of each cilium was obtained by linear interpolation from 0 to 2*π* between consecutive peaks, and the phase difference was obtained by subtracting the instantaneous phases of the two cilia. Cilia with non-persistent or irregular beating were excluded, and phase differences were calculated only for cells in which both cilia beat periodically. At a temporal resolution of 2 ms, each beat cycle was divided into 9– 12 segments, limiting the phase difference resolution to 0.5–0.7 rad.

The spatial occupancy of the cilia was quantified by summing the binary ciliary masks across all frames and dividing by the number of frames, yielding a time-averaged occupancy map. These maps were used to identify phenotypes such as partial ciliary paralysis, reduced beat amplitude and asymmetries between the two cilia.

### C. Statistical analysis

Each mutant was compared with its wild-type reference strain using a two-sided Mann–Whitney *U* test (SciPy). Significance levels are indicated as * *p <* 0.05, ** *p <* 0.01, *** *p <* 0.001 and **** *p <* 10^*−*4^; ns, not significant. No correction for multiple comparisons was applied. In box plots, boxes show the 25th and 75th percentiles, central lines show the median, and whiskers extend to the most extreme data points within 1.5 times the interquartile range. All individual data points are overlaid. Sample sizes are given in the figure legends.

## ACKNOWLEDGMENTS

We thank Pierre Gönczy for helpful discussions on this project.

## AUTHOR CONTRIBUTIONS

The research was designed by G.R.R.-S.J. and A.S. The experiments were performed by D.M.L., A.S., A.M.K. and Y.T. The data analysis was performed by A.M.K., Y.T., G.R.R.-S.J and A.S. The western blots and isolation of NFAPS were performed by G.D. The manuscript was written by G.R.R.-S.J. and edited by D.M.L., A.S., Y.T, and A.M.K.

## DATA AVAILABILITY

The data that support the plots within this paper and other findings of this study will be publicly available upon publication.

## CODE AVAILABILITY

The computer codes used in this paper are available from https://github.com/livingpatterns/chlamy_centrin_cilia

---

**Extended Data Figure 1.**
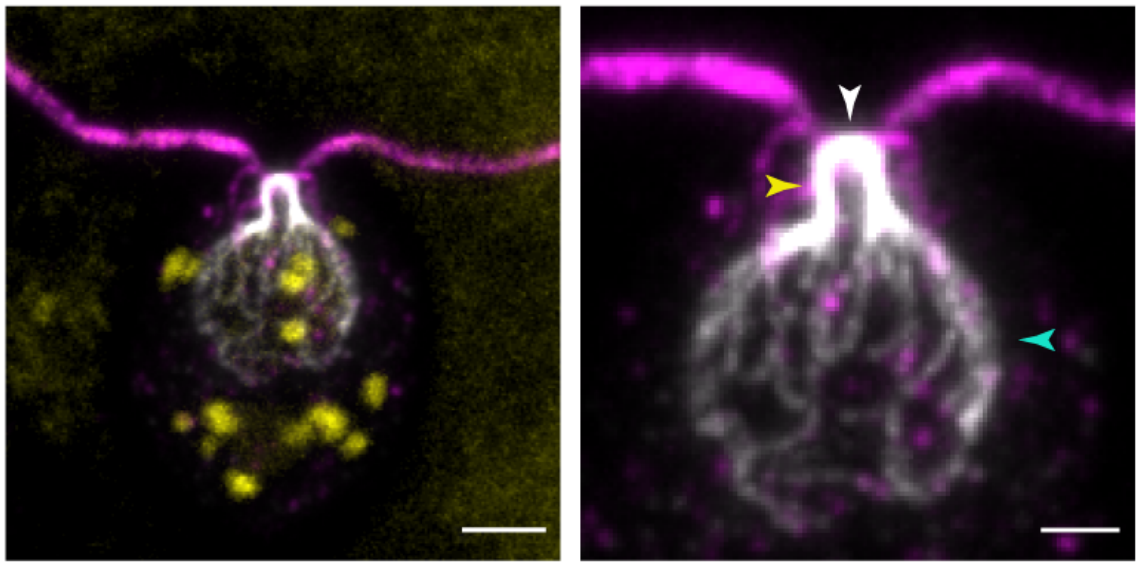
Centrin localizes to basal body-associated structures. Immunofluorescence image of *Chlamy-domonas* wild type (CC-124) cell (left) stained for centrin (white), PolyE(magenta) and DNA (yellow). Right: inset showing the centrin network at the basal body region. Centrin localization is observed at the nuclear basal body connectors (yellow arrowhead), distal striated fibers (white arrowhead) and around the nucleus as a branched network (cyan arrowhead).Scale bars: 2 µm (Left), 1 µm (Right).

**Extended Data Figure 2.**
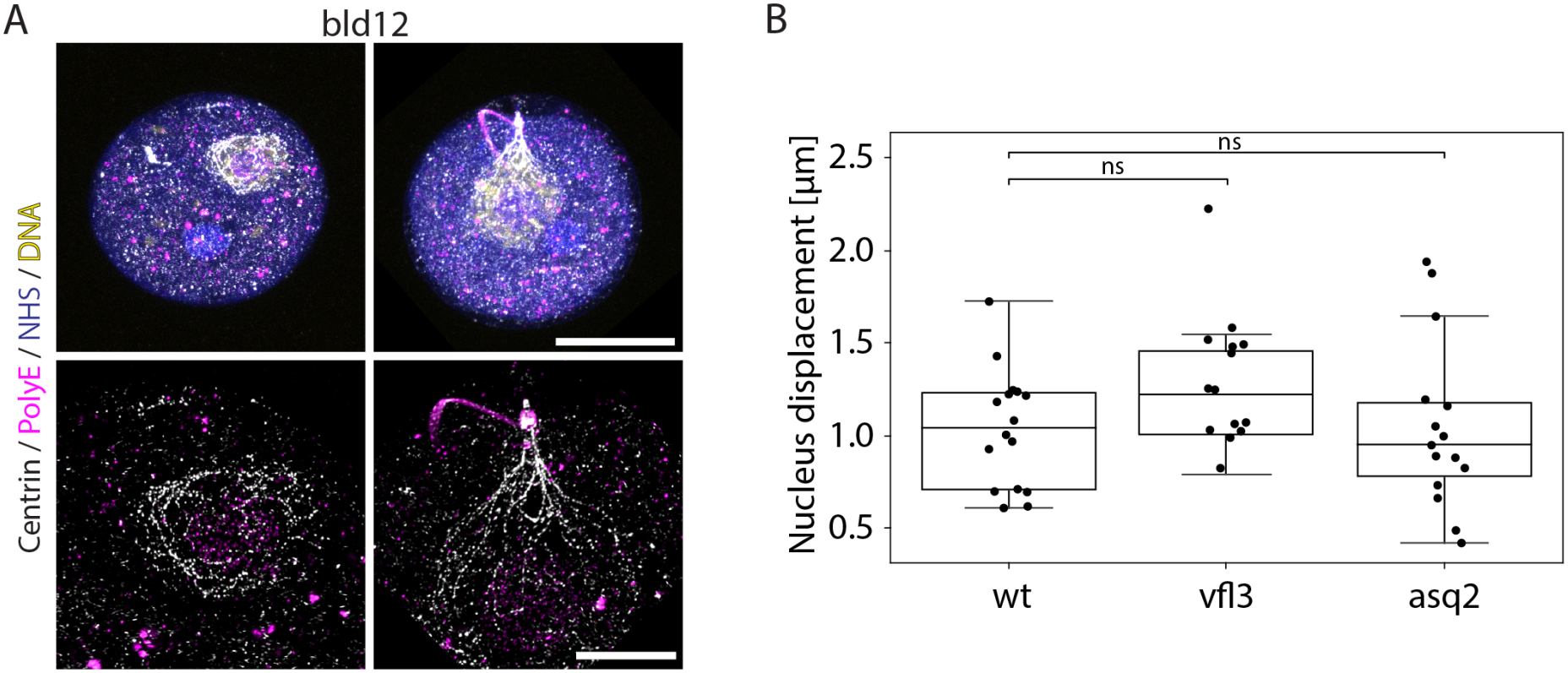
Basal bodies are required for anchoring of centrin fibres. (A) Top: Immunofluorescence images of *bld12* cells without (left) and with a basal body (right) stained for centrin (white), PolyE(magenta), NHS-ester (blue) and DNA (yellow) and imaged by ExM. Bottom: inset showing the nuclear and basal body regions for each cell (B) Nucleus displacement of wild type (CC-124), *vfl3* and *asq2* mutants from the cell centroid. Sample sizes (B) wild type (CC-124), *n*_cells_ = 16; *vfl3, n*_cells_ = 14; *asq2, n*_cells_ = 15. Scale bars: 5 µm (A, Top), 2 µm (A, Bottom).

**Extended Data Figure 3.**
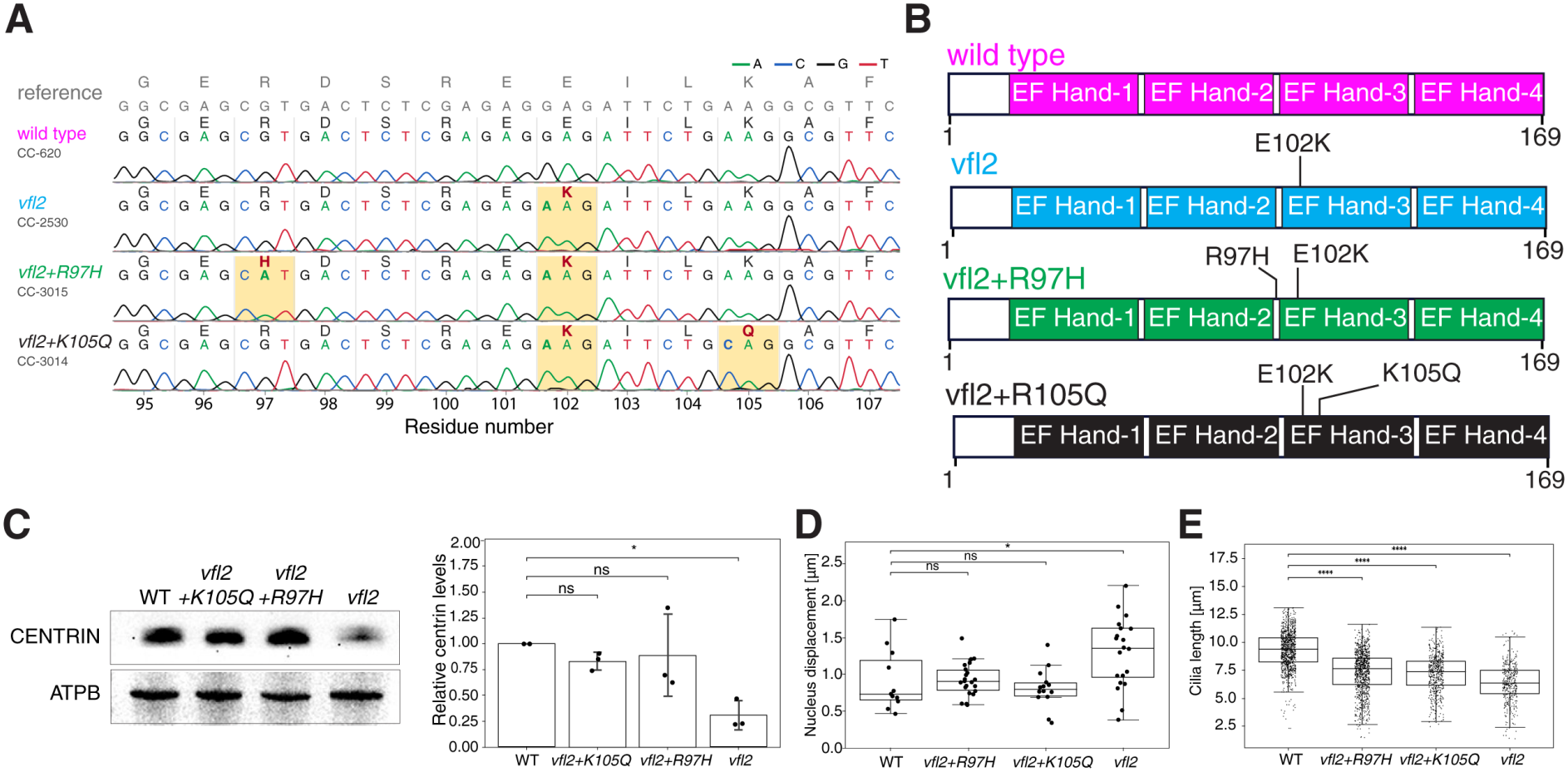
Characterization of centrin mutants. (A) Sanger sequencing data of wild type, *vfl2* and *vfl2*-derived mutants. The genomic regions surrounding the position of mutations were amplified by PCR and sequenced by Sanger sequencing. The positions of the mutations and the corresponding nucleotide and amino acid change are indicated for each mutant. (B) Schematic representation of centrin, depicting the 4 EF-Hand domains. The positions of the mutations in the strains used in this study are indicated. All the mutations are present in the third EF-Hand domain of the protein.(C) Left: Western blot of whole cell lysates of wild type (CC-620) and centrin mutants against anti-centrin antibody. Anti-AtpB is used as loading control. Right: Quantification of centrin levels in mutants relative to the wild type. The intensities of the bands corresponding to centrin signal were normalized to the intensities of corresponding AtpB signals followed by normalization of centrin signal to the wild type. Data represented are mean ± s.d from 3 independent experiments. (D) Nucleus displacement of wild type (CC-620), *vfl2+R97H, vfl2+K105Q* and *vfl2* mutants from the cell centroid. (E) Box plot of measured cilia lengths of wild type (CC-620) and centrin mutants. Sample sizes (D) wild type (CC-620), *n*_cells_ = 11; *vfl2+R97H, n*_cells_ = 22; *vfl2+K105Q, n*_cells_ = 14; *vfl2,n*_cells_ = 20 (E) wild type (CC-620), *n*_cilia_ = 1016; *vfl2+R97H, n*_cilia_ = 1001; *vfl2+K105Q, n*_cilia_ = 537; *vfl2, n*_cilia_ = 379. Box plots as in Fig. 2

**Extended Data Figure 4.**
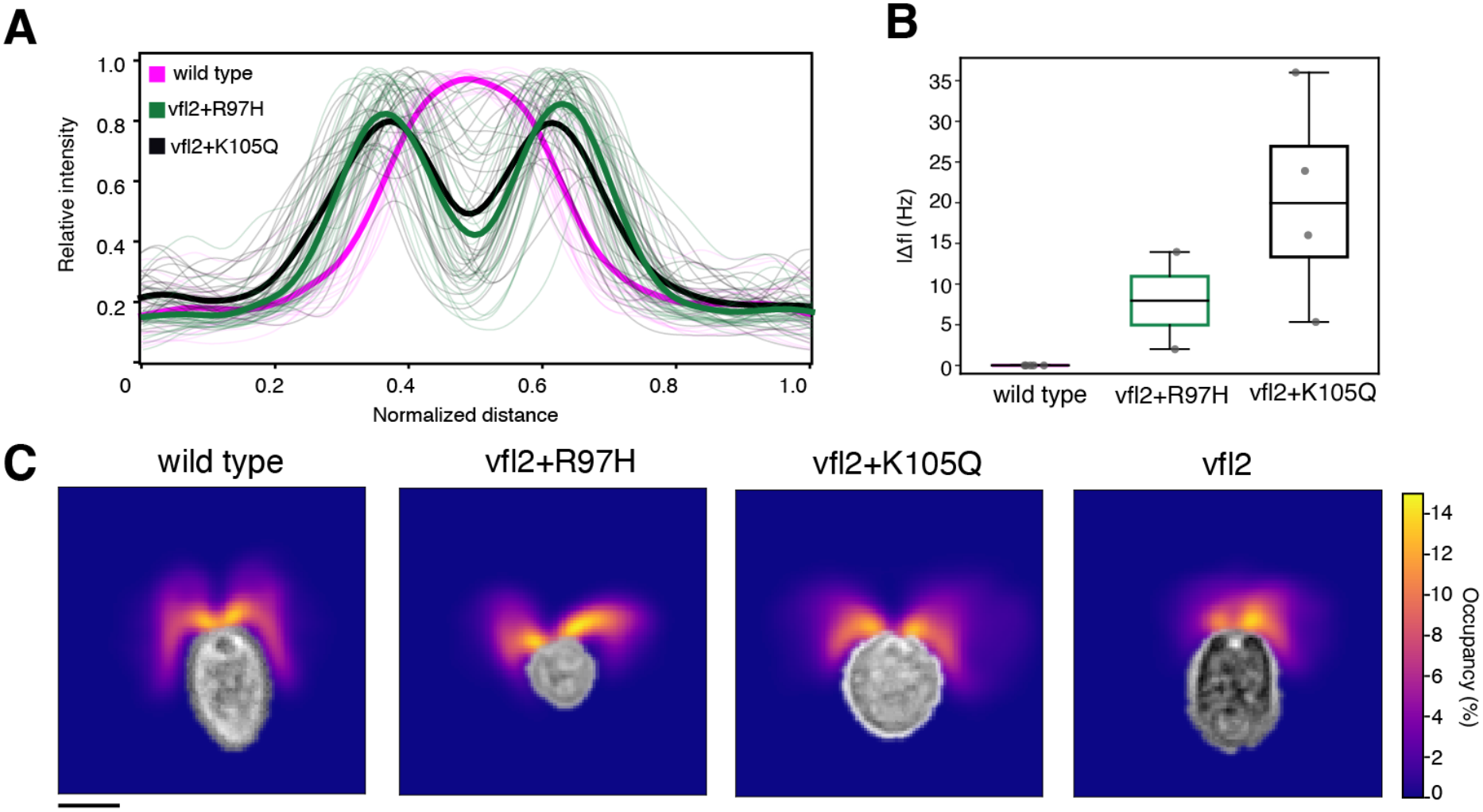
Centrin mutants show abnormal morphologies in the DSF, coupled with defective cilia coordination. (A) Fluorescence intensity profiles along the width of the DSFs in wild type, *vfl2+R97H* and *vfl2+K105Q*. Light colored curves indicate individual profiles, bold curves indicate the mean fluorescence intensities for each genotype. Sample sizes: wild type (CC-620), *n*_DSF_ = 19; *vfl2+R97H, n*_DSF_ = 25; *vfl2+K105Q, n*_DSF_ = 29. (B) Absolute difference ( |*f*_1_ − *f*_2_|) between the two cilia of individual cells across strains. Individual points represent single cells. Number of cilia pairs: wild type (CC-620), *n*_*pairs*_ = 5, *vfl2+R97H, n*_*pairs*_ = 2, *vfl2+K105Q, n*_*pairs*_ = 4. Box plots as in Fig. 2. (C) Heatmap of the ciliary spatial occupancy, showing the percentage of frames each pixel is occupied by a cilium trace, smoothed with a 2D Gaussian filter (*σ* = 3 pixels) and scaled from 0% to the maximum occupancy value. A representative example for each strain is presented. Scale bar: 5 µm

**Extended Data Figure 5.**
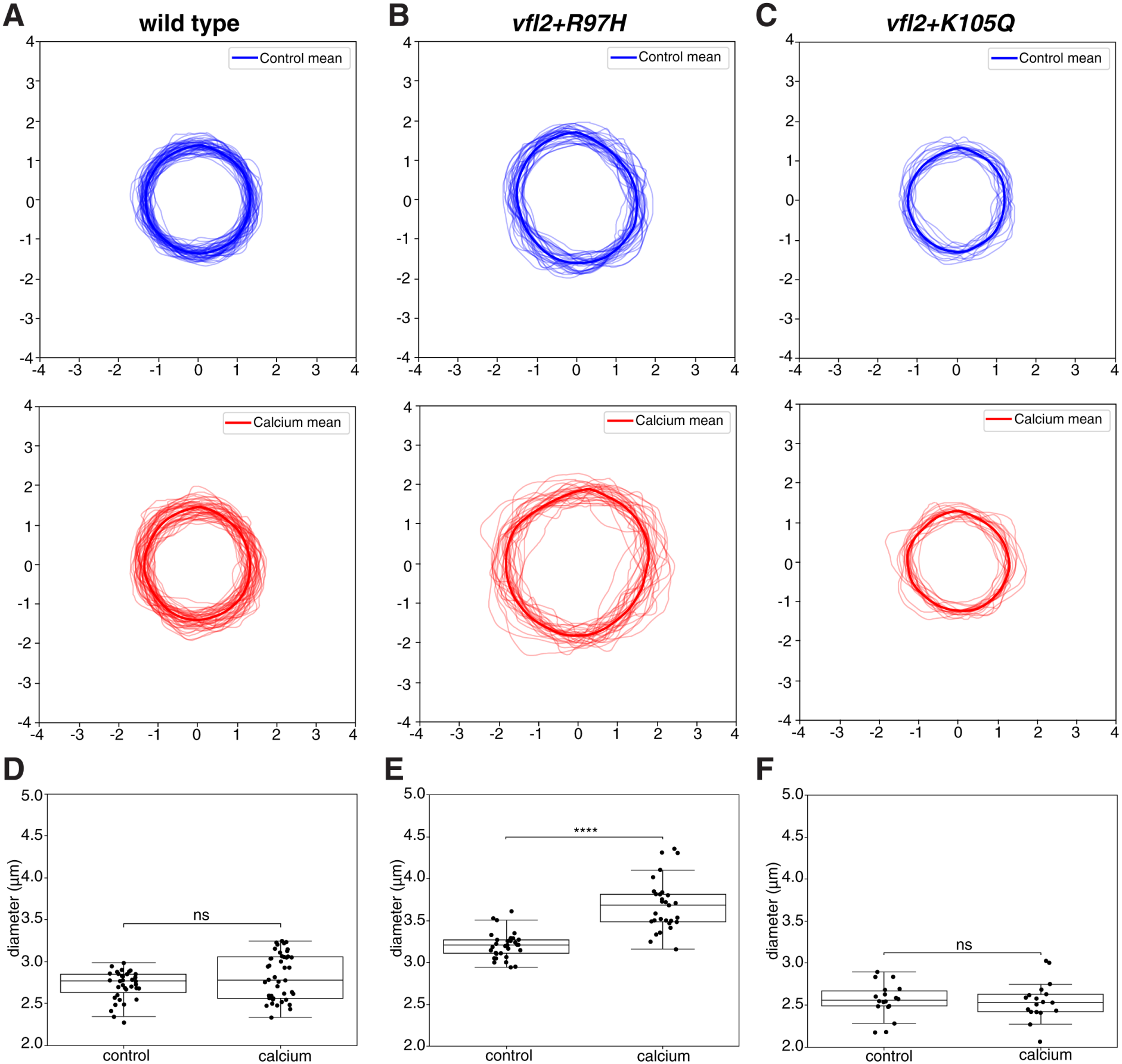
Nuclear diameter measurements in NFAPs. (A) Contours of nucleus as analyzed from immunofluorescence images of wild type NFAPs isolated without (top) and with Ca^2+^ (5 mM free Ca^2+^, bottom). Bold lines indicate the mean contour, light colored lines indicate individual contours. (B-C) As in (A) for *vfl2+R97H* (B) and *vfl2+K105Q* (C). (D) Box plot of measured nucleus diameters of wild type NFAPs isolated with and without Ca^2+^. (E-F) As in (D) for *vfl2+R97H* (E) and *vfl2+K105Q* (F). Box plots as in Fig. 2. Sample sizes (D–F): wild type (CC-620) control, *n*_nuclei_ = 12; wild type (CC-620) calcium, *n*_nuclei_ = 14;*vfl2+R97H* control, *n*_nuclei_ = 32;*vfl2+R97H* calcium, *n*_nuclei_ = 29;*vfl2+K105Q* control, *n*_nuclei_ = 18;*vfl2+K105Q* calcium, *n*_nuclei_ = 18.

**TABLE S1.** Sample sizes for all quantitative analyses. Each entry given below indicates the panel number of specific figures with the number of cells analysed per cell strain, except for Fig.3G and Fig.S3E, where the number of cilia analyzed is inidcated.

| **Panel** | **Quantity measured** | **Cell strain** | **Sample size** |
|---|---|---|---|
| Fig. 2G | Measurement of basal body distance | Wild type (cc124) | 16 |
|  |  | *Vfl3* | 13 |
|  |  | *asq2* | 15 |
| Fig. 3E | Measurement of basal body distance and angle | Wild type (cc620) | 11 |
|  |  | *Vfl2+R97H* | 22 |
|  |  | *Vfl2+K105Q* | 14 |
|  |  | *Vfl2* |  |
| Fig. 3F | Measurement of NBBC length | Wild type (cc620) | 18 |
|  |  | *Vfl2+R97H* | 25 |
|  |  | *Vfl2+K105Q* | 29 |
| Fig. 3G | Measurement of cilia length during regeneration | Wild type (cc620), 60 min | 649 |
|  |  | Wild type (cc620), 120 min | 759 |
|  |  | Wild type (cc620), 180 min | 792 |
|  |  | Wild type (cc620), 240 min | 736 |
|  |  | *Vfl2+R97H*, 60 min | 33 |
|  |  | *Vfl2+R97H*, 120 min | 103 |
|  |  | *Vfl2+R97H*, 180 min | 232 |
|  |  | *Vfl2+R97H*, 240 min | 417 |
|  |  | *Vfl2+K105Q*, 60 min | 52 |
|  |  | *Vfl2+K105Q*, 120 min | 195 |
|  |  | *Vfl2+K105Q*, 180 min | 380 |
|  |  | *Vfl2+K105Q*, 240 min | 525 |
|  |  | *Vfl2*, 60 min | 25 |
|  |  | *Vfl2*, 120 min | 109 |
|  |  | *Vfl2*, 180 min | 265 |
|  |  | *Vfl2*, 240 min | 325 |
| Fig. 3H | Cilia count | Wild type (cc620) | 807 |
|  |  | *Vfl2+R97H* | 828 |
|  |  | *Vfl2+K105Q* | 731 |
|  |  | *Vfl2* | 731 |
| Fig. 4B | DSF width | Wild type | 18 |
|  |  | *Vfl2+R97H* | 25 |
|  |  | *Vfl2+K105Q* | 29 |
| Fig. 4D | Swimming speed | Wild type | 95 |
|  |  | *Vfl2+R97H* | 44 |
|  |  | *Vfl2+K105Q* | 46 |
|  |  | *Vfl2* | 64 |
| Fig. 4E | Cilia beat frequency | Wild type | 5 |
|  |  | *Vfl2+R97H* | 5 |
|  |  | *Vfl2+K105Q* | 5 |
|  |  | *Vfl2* | 4 |
| Fig. 4H | Phase difference between 2 cilia | Wild type | 5 |
|  |  | *Vfl2+R97H* | 2 |
|  |  | *Vfl2+K105Q* | 4 |
| Fig. 5D | Measurement of Strain | Wild type | 45 |
|  |  | *Vfl2+R97H* | 29 |
|  |  | *Vfl2+K105Q* | 17 |
| Extended data Fig. S2B | Measurement of nucleus displacement | Wild type(CC-124) | 16 |
|  |  | *vfl3* | 14 |
|  |  | *asq2* | 15 |
| Extended data Fig. S3D | Measurement of nucleus displacement | Wild type(CC-620) | 11 |
|  |  | *vfl2+R97H* | 22 |
|  |  | *vfl2+K105Q* | 14 |
|  |  | *vfl2* | 20 |
| Extended data Fig. S3E | Measurement of cilia length | Wild type(CC-620) | 1016 |
|  |  | *vfl2+R97H* | 1001 |
|  |  | *vfl2+K105Q* | 537 |
|  |  | *vfl2* | 379 |
| Extended data Fig. S4A | Fluorescence intensity profiles | Wild type(CC-620) | 19 |
|  |  | *vfl2+R97H* | 25 |
|  |  | *vfl2+K105Q* | 29 |
| Extended data Fig. S4B | Absolute ciliary beat frequency difference | Wild type(CC-620) | 5 |
|  |  | *vfl2+R97H* | 2 |
|  |  | *vfl2+K105Q* | 4 |
| Extended data Fig. S5D | Measurement of nucleus diameter | Wild type(CC-620) control | 12 |
|  |  | Wild Type calcium | 14 |
| Extended data Fig. S5E | Measurement of nucleus diameter | *vfl2+R97H* control | 32 |
|  |  | *vfl2+R97H* calcium | 29 |
| Extended data Fig. S5F | Measurement of nucleus diameter | *vfl2+K105Q* control | 18 |
|  |  | *vfl2+K105Q* calcium | 18 |

Movie S1. **Ciliary beating in free-swimming wild-type and centrin mutant cells**. High-speed DIC recordings of three representative cells (rows) each of wild type (CC-620), *vfl2* +R97H, *vfl2* +K105Q and *vfl2* (columns). Recordings were acquired at 1000 fps, registered to correct for cell motion and rotated so that the anterior end of the cell points upward (black corners result from this registration). They are shown at 500 fps, with 2 ms between frames, and played back at 30 fps. Time stamps indicate elapsed time. In wild-type cells, the two cilia beat in synchrony with a symmetric breaststroke. In *vfl2* +R97H cells, one cilium frequently beats defectively, switching between rigid paralysis and slow, broad strokes. In *vfl2* +K105Q cells, the two cilia beat at different frequencies and drift in and out of phase. *vfl2* cells show the most severe phenotype, with irregular, aperiodic beating. Scale bar: 10 µm.

